# MIF-mediated reprogramming of myeloid lineage within the glioma tumor microenvironment impacts the efficacy of immune stimulatory gene therapy

**DOI:** 10.64898/2026.01.13.699283

**Authors:** Ziwen Zhu, Noah C. Kanis, Anthony E. George, Michael Albdewi, Yingxiang Li, Anzar A. Mujeeb, Brandon L. McClellan, Gurveer Singh, Jialin Liu, Wajd N. Al-Holou, Jason A. Heth, Joshua D. Welch, Justin D. Lathia, Pedro R. Lowenstein, Richard Bucala, Maria G. Castro

## Abstract

Gliomas with mutant isocitrate dehydrogenase 1 (mIDH1) represent a distinct subgroup of brain tumors characterized by unique metabolic and immunological profiles compared to wildtype IDH1 (wtIDH1) gliomas. Despite recent progress, the cellular mechanisms underlying tumor progression and immune modulation in these subtypes remain poorly understood. In this study, we employed single-cell RNA sequencing (scRNA-seq) to characterize the cellular heterogeneity of wtIDH1 and mIDH1 gliomas, with a particular focus on myeloid cell populations. Our analyses revealed a marked reduction of monocyte-derived tumor-associated macrophages (Mo-TAMs) and lower expression of macrophage migration inhibitory factor (MIF) in mIDH1 gliomas, which was attributable to epigenetic reprogramming. Mechanistic studies using MIF and CD74 knockout mice demonstrated that the MIF-CD74 axis plays a crucial role in regulating the glioma immune microenvironment, thereby driving tumor growth and progression. Importantly, the combination of immune-stimulatory gene therapy (HSV1-thymidine kinase/Fms-like tyrosine kinase 3 ligand; TK/Flt3L) with MIF inhibition significantly extended survival in models of wtIDH1 glioma. These findings highlight the therapeutic potential of targeting the MIF-CD74 pathway and underscore the importance of integrating immunomodulatory strategies for the treatment of glioma.

**Highlights:**

- Mutant IDH1 gliomas exhibit fewer Mo-TAMs and increased Mg-TAMs
- Mutant IDH1 gliomas have less MIF expression via epigenetic reprogramming.
- Mesenchymal wtIDH1 glioma cells are main source of MIF.
- MIF inhibition plus immune stimulatory gene therapy extends survival wtIDH1 glioma.

## Introduction

Isocitrate dehydrogenase (IDH) mutations are hallmark genetic lesions in gliomas, fundamentally altering tumor biology and patient outcomes. Approximately 90% of IDH1 mutations occur at exon 4, codon 132, resulting in a substitution of arginine by histidine (R132H), while less frequent mutations are found at IDH2 codon R172. These hotspot mutations, notably common in astrocytomas, are often accompanied by co-occurring alterations in ATRX and TP53, further shaping tumor pathogenesis^1,2^. The mutant IDH1 enzyme acquires neomorphic activity, catalyzing the conversion of α-ketoglutarate (αKG) to the oncometabolite 2-hydroxyglutarate (2-HG) ^3^.Accumulation of 2-HG inhibits key epigenetic regulators, including histone demethylases (KDMs) and DNA demethylases (TET2), forcing a global hypermethylation phenotype and extensive transcriptomic reprogramming^4,5^.

Recent studies highlight a central role for mutant IDH1 (mIDH1) in sculpting the immunological landscape of the glioma tumor microenvironment (TME)^6,7^. The TME of gliomas is predominantly infiltrated by myeloid cells, especially tumor-associated macrophages (TAMs) and myeloid-derived suppressor cells (MDSCs), which orchestrate both tumor-promoting processes and immune evasion^7–9^. TAMs arise from two major sources: peripheral monocytes (monocyte-derived TAMs, or Mo-TAMs), and tissue-resident microglia (microglia-derived TAMs, or Mg-TAMs)^7,10^. Whereas homeostatic monocytes perform protective functions in healthy tissue, those conscripted into the glioma TME acquire tumor-supportive roles, distinguished by gene expression signatures, a preference for the tumor core, and elevated abundance in wildtype IDH1 (wtIDH1) and recurrent tumors ^7,8,10^.TAMs directly interact with neoplastic cells, facilitate tumor progression, and establish a profoundly immunosuppressive microenvironment.

Although several mIDH1-dependent aspects of the glioma immune response have been described, many mechanisms underlying TME immune function remain insufficiently characterized. In this study, we systematically characterize the immune TME of human and murine wtIDH1 and mIDH1 gliomas, focusing on myeloid cell diversity using single-cell RNA sequencing (scRNA-seq). Our analysis reveals a marked reduction in monocyte-derived TAMs and significantly lower expression of macrophage migration inhibitory factor (MIF) in mIDH1 gliomas, a difference attributable to epigenetic reprogramming. Through mechanistic studies using MIF- and CD74-knockout mouse models, we identify the MIF-CD74 signaling axis, which is deficient in mIDH1 glioma, as a key regulator of immune cell composition, tumor progression, and survival.

To explore therapeutic opportunities, we evaluated immune-stimulatory gene therapy combining HSV1-thymidine kinase and Fms-like tyrosine kinase 3 ligand (TK/Flt3L), a strategy developed in our laboratory that has completed a phase I clinical trial in adult high-grade glioma patients (ClinicalTrials.gov identifier: NCT01811992)^11^. Importantly, combining TK/Flt3L gene therapy with pharmacological MIF inhibition substantially extends survival in genetically engineered, immunocompetent wtIDH1 glioma models.

Collectively, our findings elucidate the critical impact of mIDH1 on the phenotypic and functional reprogramming of myeloid cells in the glioma microenvironment and support the integration of MIF-CD74 pathway inhibition into immunomodulatory treatment strategies. This work underscores the translational potential of targeting the MIF-CD74 pathway in combination with gene therapy to improve therapeutic outcomes for glioma patients.

## Result

### Single cell RNA sequencing identified MIF pathway dysregulation in mIDH1 glioma

We analyzed single-cell RNA sequencing (scRNA-seq) data from 35 human glioma cases to characterize the composition of the tumor microenvironment (TME) (Figure 1A). Specimens included tumors from patients with wild-type IDH1 (wtIDH1) glioma (n = 25) and mutant IDH1 (mIDH1) astrocytoma. Myeloid cells were further classified into monocytic MDSCs, monocyte-derived TAMs (Mo-TAMs), microglia-derived TAMs (Mg-TAMs), and dendritic cells (DCs), while glioma cells were categorized into astrocyte-like (AC-like), oligodendrocyte progenitor cell-like (OPC-like), neural progenitor cell-like (NPC-like), and mesenchymal-like (MES-like) populations according to GBmap annotation^12^. Notably, mIDH1 astrocytoma samples exhibited an enrichment of OPC-like cells and a reduction in MES-like cells (Figure 2B).

**Figure 1.**
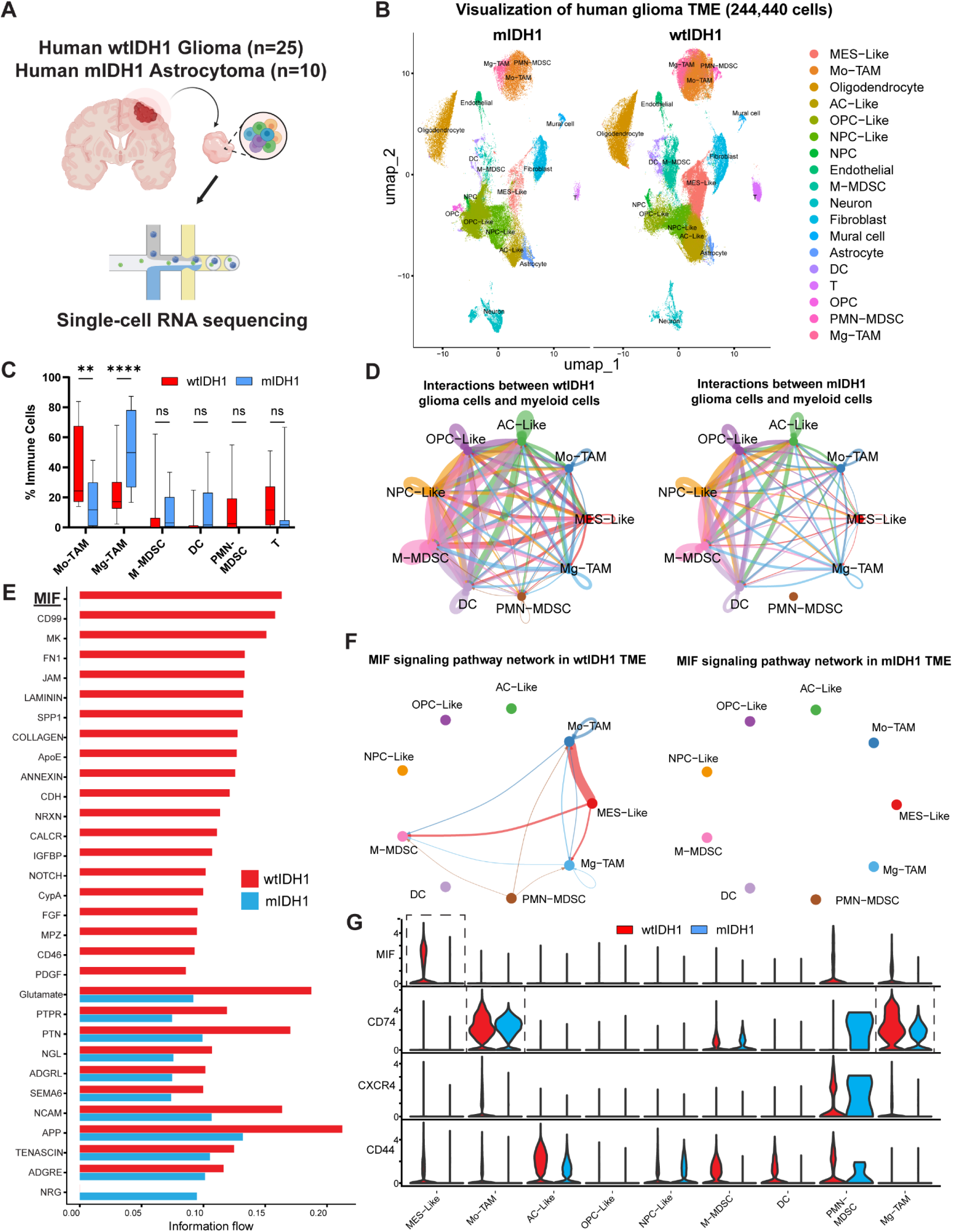
Single cell RNA sequencing identified MIF pathway dysregulation in mIDH1 glioma. **(A)** Illustration of human glioma specimens in our scRNA-seq dataset. **(B)** UMAP embedding of wtIDH1 and mIDH1 scRNA-seq. **(C)** Immune cell percentages from wtIDH1 and mIDH1 samples. **(D)** Tumor-immune cell communication in wtIDH1 and mIDH1 glioma analyzed by CellChat. **(E)** Difference of signaling pathways in wtIDH1 and mIDH1 glioma. **(F)** Circle plot depicting the MIF-CD74 crosstalk in wtIDH1 and mIDH1 glioma. **(G)** Violin plot depicting the MIF-CD74 signaling pathway related gene expression in wtIDH1 and mIDH1 glioma.

**Figure 2.**
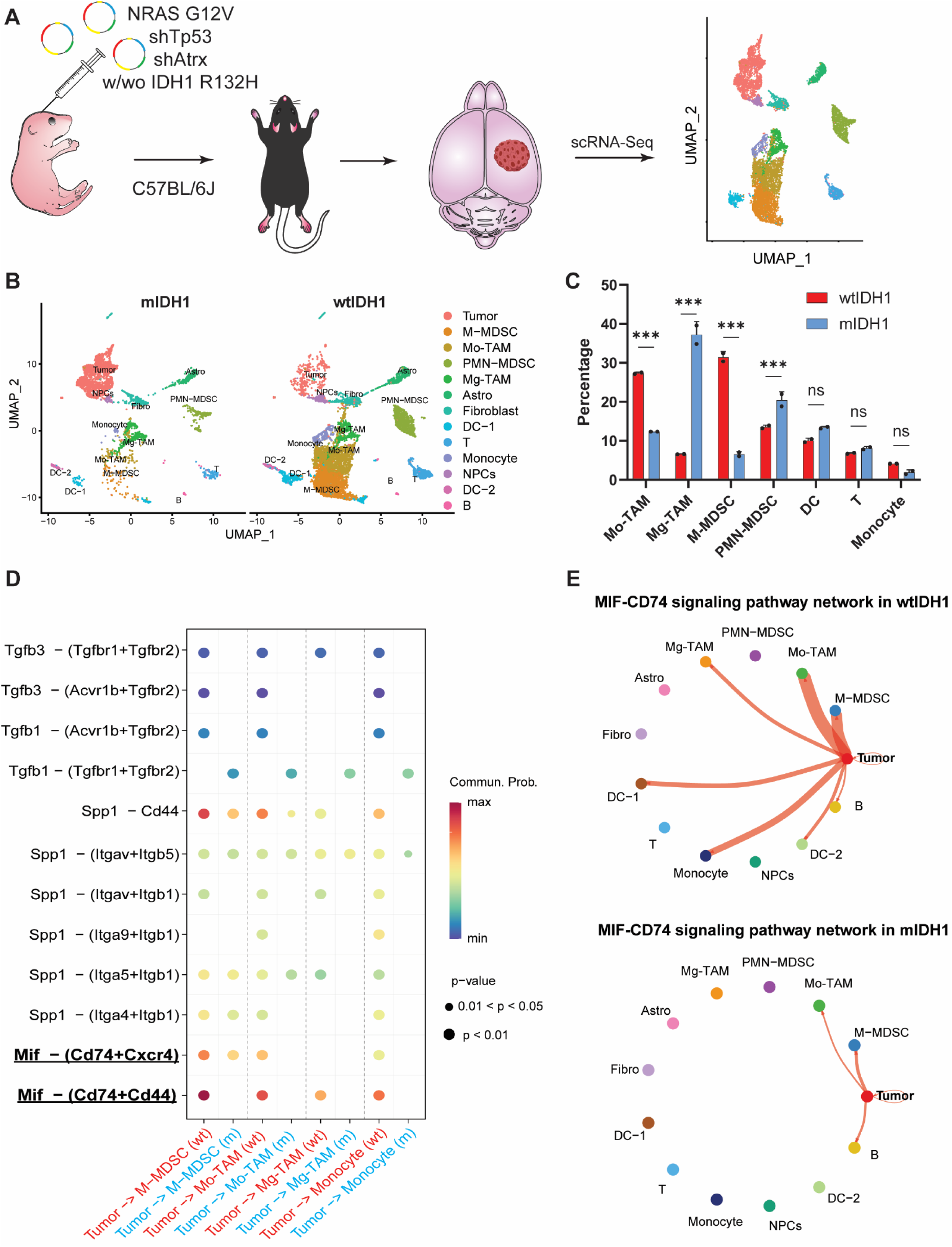
Development of immunocompetent mouse wtIDH1 and mIDH1 glioma. **(A)** Illustration of experimental procedure followed for the scRNA-seq characterization of wtIDH1 and mIDH1 mouse TME. **(B)** UMAP embedding of wtIDH1 and mIDH1 scRNA-seq. **(C)** Immune cell percentages from wtIDH1 and mIDH1 samples. **(D)** Tumor-immune cell communication in wtIDH1 and mIDH1 glioma analyzed by CellChat. **(E)** Circle plot depicting the MIF-CD74 crosstalk in wtIDH1 and mIDH1 glioma.

Analysis of TIME composition revealed a higher proportion of Mo-TAMs in wtIDH1 glioma, whereas Mg-TAMs were more prevalent in mIDH1 astrocytoma (Figure 1C). To investigate intercellular communication within the TME, we performed CellChat analysis, which demonstrated a significant decrease in interactions among MES-like cells, Mo-TAMs, and Mg-TAMs in mIDH1 astrocytoma compared to wtIDH1 glioma (Figure 1D).

Further evaluation of 3,233 signaling pathways using the CellChat database identified 20 pathways significantly enriched in wtIDH1 glioma (Figure 1E), with the MIF pathway showing the most pronounced difference. Detailed analysis revealed that MIF is predominantly secreted by MES-like wtIDH1 glioma cells and acts on Mo-TAMs but is nearly absent in the mIDH1 TME (Figure 1F). Violin plots confirmed high MIF expression in MES-like wtIDH1 glioma cells, while CD74 was mainly expressed on Mo-TAMs and Mg-TAMs (Figure 1G).

To further validate MIF pathway dysregulation in mIDH1 glioma, we developed an immunocompetent mouse model using the Sleeping Beauty (SB) transposon system, which enables de novo induction of brain tumors through non-viral, transposon-mediated integration of plasmid DNA into subventricular zone cells of neonatal mice. Given that RAS/MAPK pathway activation is associated with MES-like glioma cells^13–15^—the principal source of MIF production (Figure 1F)—we modeled wtIDH1 and mIDH1 gliomas by delivering plasmids encoding NRAS G12V, short hairpin RNAs targeting Trp53 and Atrx (NPA), and, for mIDH1 gliomas, an additional IDH1 R132H plasmid (NPAI). To characterize cellular crosstalk and immune infiltration in the tumor microenvironment, we performed scRNA-seq on SB-induced wtIDH1 and mIDH1 gliomas (Figure 2A, Figure S18). Immune cell analysis revealed a significant reduction in monocytic myeloid-derived suppressor cells (M-MDSCs) and monocyte-derived TAMs (Mo-TAMs), accompanied by a relative increase in microglia-derived TAMs (Mg-TAMs) in mIDH1 gliomas, mirroring observations from human glioma datasets (Figure 2B-C).

Furthermore, wtIDH1 gliomas displayed enrichment of MES-like cells and a reduction in OPC-like cells compared to mIDH1 gliomas (Figure S2B). CellChat analysis of tumor–myeloid cell interactions revealed a significant dysregulation of the MIF-CD74 signaling pathway in mIDH1 gliomas, consistent with findings in human samples (Figure 2D-E).

To further determine whether MIF expression varies with IDH1 mutation status in glioma patients, we analyzed publicly available RNA-seq data from The Cancer Genome Atlas (TCGA) (Figure S3A). Our analysis revealed a significant increase in log2-normalized MIF expression in wtIDH1 gliomas compared to mIDH1 gliomas. Stratifying patients by MIF expression demonstrated that those with high MIF levels in the wtIDH1 group experienced poorer overall survival compared to those with low MIF expression (Figure S3B). In contrast, MIF expression was not associated with survival outcomes within the mIDH1 glioma cohort. This notable difference in overall survival suggests that a subset of wtIDH1 gliomas with elevated MIF expression is associated with unfavorable clinical prognosis. Additional analysis using data from the Chinese Glioma Genome Atlas (CGGA) corroborated these findings, showing results consistent with those from TCGA (Figure S3C-D).

### MIF expression is epigenetically downregulated in mIDH1 glioma cells

To validate our scRNA-seq findings regarding differential MIF expression in wtIDH1 versus mIDH1 glioma cells, We first implanted wtIDH1 and mIDH1 glioma NS into MIF WT mice (Figure 4A). Consistent with patient survival outcomes, mice bearing mIDH1 gliomas exhibited prolonged survival compared to those with wtIDH1 gliomas (Figure 4B). ELISA analysis of serum showed significantly lower MIF levels in mIDH1 glioma-bearing mice (Figure 4C). We also performed western blot and ELISA assays to quantify MIF levels in both mouse and human glioma cells (Figure S4A). Western blot analysis revealed that mouse mIDH1 glioma cells exhibited decreased MIF expression relative to wtIDH1 controls (Figure S4B), with lower MIF concentrations detected in the conditioned media as well (Figure S4C). To confirm the results from mouse NS, we also used human glioma cells, SJ-GBM2-wtIDH1 and SJ-GBM2-mIDH1, where mIDH1 cells demonstrated reduced MIF expression (Figure S4D) and diminished MIF secretion into the conditioned media (Figure S4E).

Mutation of IDH1 in glioma cells leads to the production of the oncometabolite 2-HG, which inhibits DNA hydroxylases and histone demethylases, ultimately driving widespread epigenetic reprogramming. We hypothesized that the observed reduction in MIF expression is a consequence of mIDH1-induced epigenetic alterations. To test this, we conducted Cleavage Under Targets and Release Using Nuclease (CUT&RUN) analysis to evaluate enrichment of the active histone mark H3K4me3 (Figure 3D).

**Figure 3.**
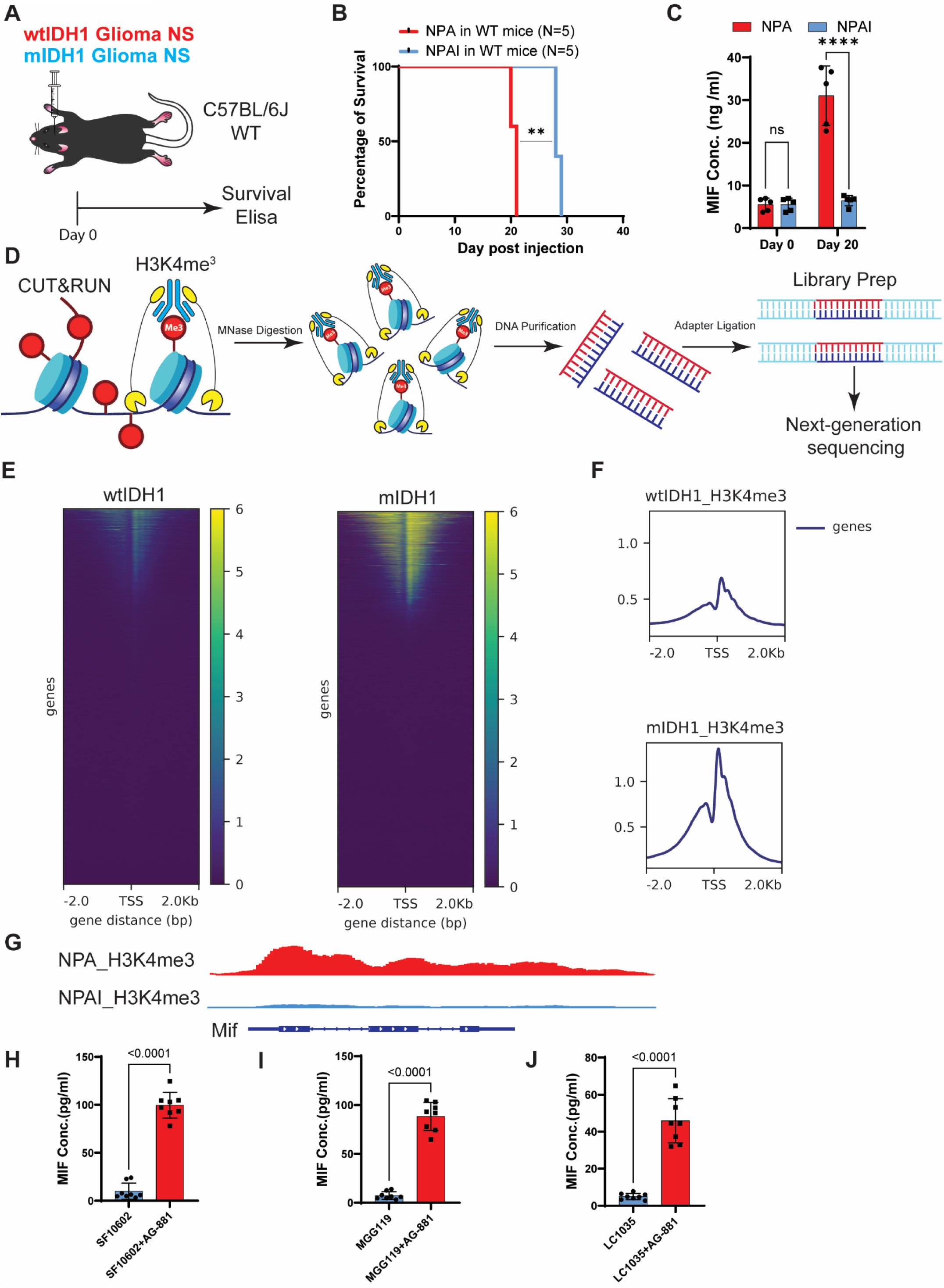
MIF expression is epigenetically downregulated in mIDH1 glioma cells. **(A)** Illustration of experimental procedure followed for the survival study and Elisa in WT mice implanted with wtIDH1 and mIDH1 mouse cells. **(B)** Kaplan-Meier survival curve shows the survival of WT mice implanted with wtIDH1 and mIDH1 mouse cells. Statistical significance was assessed using the Log-rank (Mantel-Cox) test. **(C)** Serum concentration of MIF analyzed by ELISA in the WT mice implanted with wtIDH1 and mIDH1 mouse cells. **(D)** Experimental outline for the Cleavage under targets and release using nuclease (CUT&RUN) analysis. Three biological replicates of wtIDH1 and mIDH1 neurospheres (NS) were used to perform the CUT&RUN. **(E)** Heat maps showing and H3K4me3 differential peaks ± 2 kilo–base (Kb) pair from the gene center. Each row represents a distinct gene. **(F)** Density plots of CUT&RUN data. Average density signal is shown for 2 kb regions surrounding transcription start sites. **(G)** Occupancy of H3K4me3 on the Mif in wtIDH1 and mIDH1 mouse cells. **(H-J)** The concentration of MIF in the human patient derived mIDH1 cell culture conditioned media treated with DMSO and mIDH1 inhibitor was analyzed by ELISA.

Heatmaps centered on peak midpoints revealed significant changes in histone mark deposition in mIDH1 neurospheres (Figure 3E-F), with a pronounced decrease in H3K4me3 enrichment at the MIF promoter (Figure 4G). To further confirm the epigenetic regulation of MIF by mIDH1, we treated patient-derived mIDH1 glioma cells (SF10602, MGG119 and LC1035) with either vehicle or 50 nM of the mIDH1 inhibitor AG-881 for three days, a treatment known to reverse mIDH1-mediated epigenetic changes.

Following inhibitor treatment, we observed a significant increase in MIF secretion in the conditioned media (Figure 4H-J). Taken together, these results demonstrate that mIDH1 exerts epigenetic control over MIF expression in glioma cells.

### Glioma cells are the main source of MIF

Since previous studies and our scRNA-seq data indicate that macrophages, including TAMs, can also secrete MIF and exhibit self-activation of the MIF pathway, we sought to determine the predominant origin of MIF in glioma. To assess the contribution of host-derived MIF, wtIDH1 glioma NS were implanted into both MIF WT and MIF knockout (MIF-/-) mice (Figure S5A); survival rates were similar between the groups, indicating that host MIF does not significantly influence glioma progression (Figure S5B). End-stage serum MIF concentrations were also comparable between MIF WT and MIF-/-mice and were approximately tenfold higher than baseline (Figure S5C). Similar experiments were conducted with mIDH1 glioma NS implanted in MIF WT and MIF-/-mice (Figure S5D), again showing no survival difference (Figure S5E) and no significant variation in serum MIF levels between groups, which were also similar to baseline (Figure S5F). These collective results demonstrate that wtIDH1 glioma cells, rather than host-derived cells, are the predominant source of MIF in the glioma microenvironment.

### MIF knockout in glioma delays tumor progression and regulates myeloid cells in the TME

Recognizing the elevated MIF expression in wtIDH1 gliomas and the predominant role of glioma cells as the source of MIF, we investigated the impact of tumor-derived MIF on survival by generating MIF knockout (KO) wtIDH1 genetically engineered mouse models (GEMMs) using the Sleeping Beauty (SB) transposase system. Spontaneous gliomas were induced in both wild-type and MIF-/- mice by expressing NRAS alongside shRNAs targeting Trp53 and Atrx (Figure 4A). Tumors developed in both genotypes (NPA MIF+/+ and NPA MIF-/-), with the NPA MIF-/- group (MS=134) exhibiting longer median survival compared to MIF wildtype controls (MS=79) (Figure 4B). Western blot analysis confirmed effective MIF knockout in isolated and cultured neurospheres (Figure 4C), and in vitro cell proliferation assays showed no significant differences, suggesting that MIF primarily influences the immune microenvironment rather than intrinsic tumor growth (Figure 4D).

To assess patterns of immune infiltration and elucidate the role of MIF within the TME, we intracranially implanted NPA MIF+/+ and NPA MIF-/- cells into C57BL/6 mice. Upon onset of symptoms, tumors were harvested for flow cytometry (Figure 4E). NPA MIF-/-glioma-bearing mice demonstrated significantly prolonged survival compared to their NPA MIF+/+ counterparts, indicating a direct effect of tumor cell-derived MIF on host outcome (Figure 4E). End-stage serum MIF levels in NPA MIF-/- glioma-bearing mice were comparable to baseline, further confirming glioma cells as the primary source of circulating MIF (Figure 4F).

Flow cytometric analysis revealed a complete reduction in monocyte-derived macrophages (CD45+ CD11b+ F4/80+) and a substantial decrease in dendritic cells (CD45+ CD11b+ CD11c+), from 40% to 15%, within NPA MIF-/- gliomas (Figures 4G-J). Analysis of myeloid-derived suppressor cells (MDSCs) showed similar proportions of M-MDSCs (CD45+ CD11b+ Ly6Chi Ly6G-) and PMN-MDSCs (CD45+ CD11b+ Ly6Clo Ly6G+) in both NPA MIF+/+ and NPA MIF-/- tumors (Figure S5A-B). T cell infiltration, as measured by CD45+/CD3+ populations, remained unchanged between groups (Figure S5C). Collectively, these results suggest that MIF orchestrates the recruitment and maintenance of mature myeloid cells within the glioma TME, thereby influencing tumor progression and host survival.

**Figure 4.**
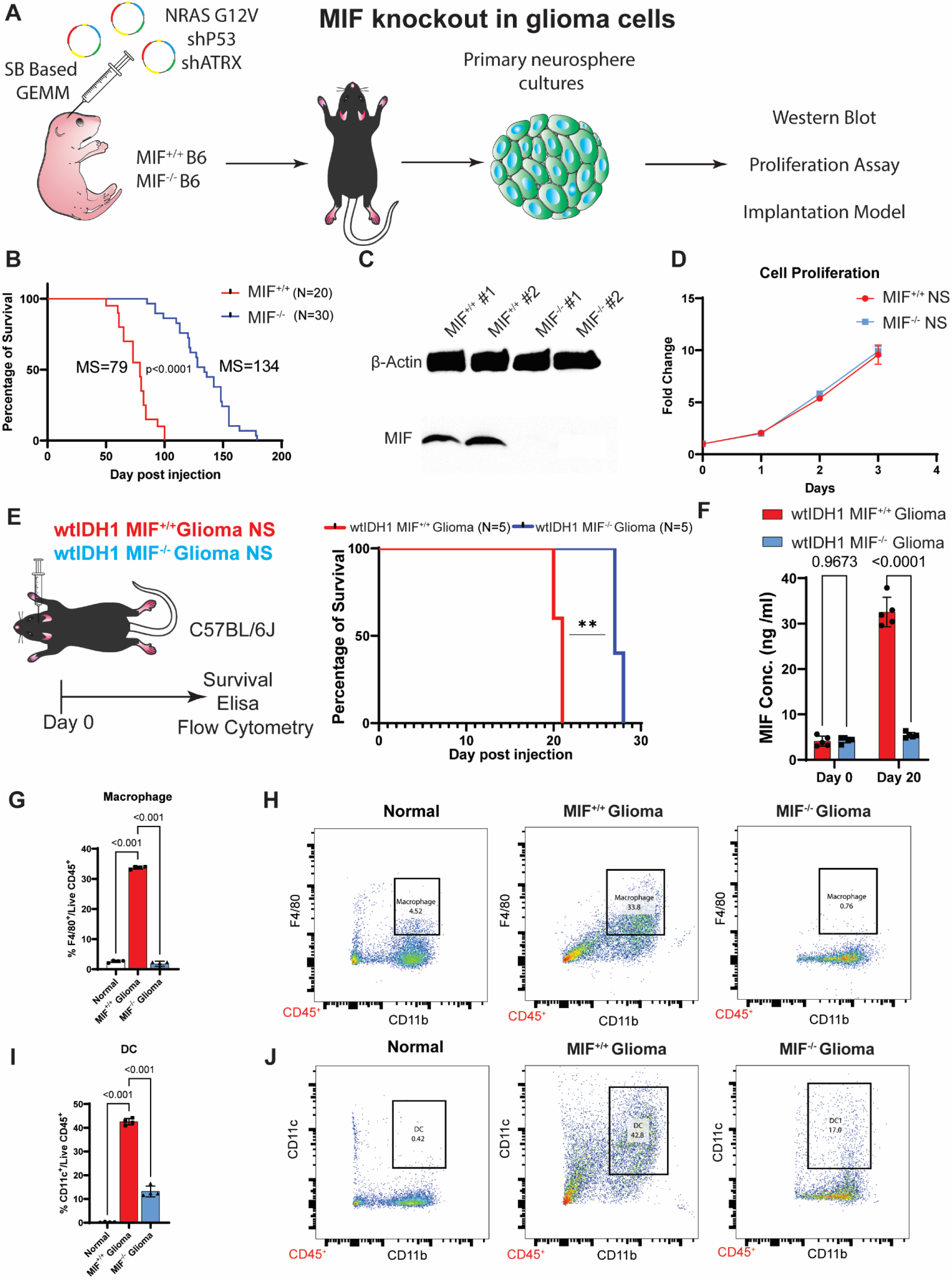
MIF knockout in glioma delays tumor progression and regulates myeloid cells in the TME. **(A)** Illustration of experimental procedure followed for the development of MIF^-/-^ glioma model using sleeping beauty transpose. **(B)** Kaplan-Meier survival curve for SB-derived MIF-WT (n=20) or MIF^-/-^ (n=30) tumor bearing animals. **** p<0.0001, Mantel-Cox test. **(C)** Western blot of MIF in MIF-WT and MIF^-/-^ mouse glioma cells. **(D)** Proliferation of MIF-WT and MIF^-/-^ mouse glioma cells *in vitro*. **(E)** Illustration of experimental procedure followed for the survival study and Elisa in WT mice implanted with MIF WT and MIF^-/-^ mouse glioma cells. **(F)** Serum concentration of MIF analyzed by ELISA in WT mice implanted with MIF-WT and MIF^-/-^ mouse glioma cells. **(G-H)** The total proportion of macrophages is shown as the percentage of CD45+/CD11b+/F4/80+ cells. **(I-J)** The total proportion of DC is shown as the percentage of CD45+/CD11b+/CD11c+.

### Knockout of CD74 in host delays tumor progression and regulates myeloid cells in the TME

Given that MIF exerts its cytokine function by binding to CD74 on immune cells—subsequently recruiting the co-receptor CD44 and engaging CXCR4, we investigated the influence of CD74 on the glioma TIME. To assess the role of CD74, we implanted NPA MIF+/+ cells into both wildtype and CD74 knockout (CD74-/-) mice (Figure 5A). CD74-/- mice exhibited significantly prolonged survival compared to wildtype controls (Figure 5B). At symptomatic stages, tumors were extracted and analyzed by flow cytometry (Figure 5A). Quantification of myeloid-derived suppressor cell populations revealed similar proportions of M-MDSCs and PMN-MDSCs in both genotypes (Figure 5C-D). However, flow cytometric analysis demonstrated a marked reduction in monocyte-derived macrophages (CD45+ CD11b+ F4/80+), from 35% to 20%, and in dendritic cells (CD45+ CD11b+ CD11c+), from 45% to 25%, within tumors implanted in CD74-/- mice (Figures 5E-H).

**Figure 5.**
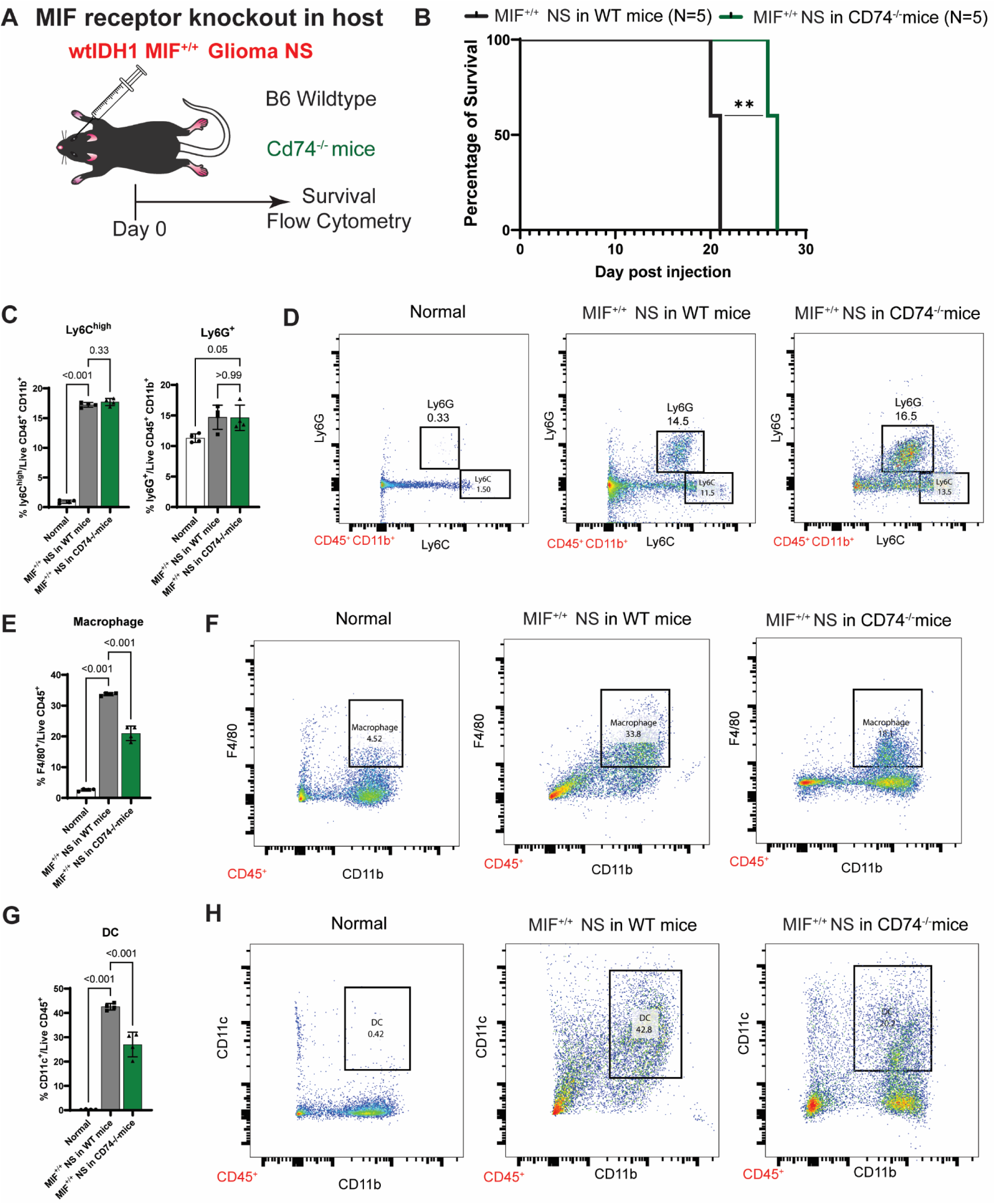
Knockout of CD74 in host delays tumor progression and regulates myeloid cells in the TME. **(A)** Illustration of experimental procedure followed for the characterization of immune cells in WT and CD74^-/-^ mice implanted with WT mouse glioma cells. **(A)** Kaplan-Meier survival curve shows the survival of WT and CD74^-/-^ mice implanted with WT mouse glioma cells. **(C-D)** The proportion of cells with Monocytic (CD45+/CD11b+/Ly6G-Ly6Chi) or Polymorphonuclear (CD45+/CD11b+/Ly6G+Ly6Clo) myeloid derived suppressor cell. **(E-F)** The total proportion of macrophages is shown as the percentage of CD45+/CD11b+/F4/80+ cells. **(G-H)** The total proportion of DC is shown as the percentage of CD45+/CD11b+/CD11c+.

To further delineate the cell-specific effects of CD74, we crossed LysM-cre mice with CD74 floxed counterparts to selectively delete CD74 in myeloid lineages (Figure S7A). Consistent with global CD74 knockout, these conditional mutants displayed similar improvements in survival (Fig. S7B) and reductions in macrophage and dendritic cell infiltration (Fig. S7C-H). Taken together, these findings demonstrate that the MIF-CD74 axis is a key regulator of mature myeloid cell composition within the glioma TME, significantly impacting tumor progression and survival.

### MIF inhibition enhances efficacy of immune stimulatory gene therapy in wtIDH1 glioma

As we found MIF signaling pathway regulates the immune microenvironment in glioma, we evaluated the efficacy of a cytotoxic immune-stimulatory TK/Flt3L gene therapy combined with MIF inhibition. This gene therapy entails injecting viruses encoding Thymidine kinase (TK) and FMS-like tyrosine kinase 3 ligand (Flt3L) into the tumor, followed by ganciclovir (GCV) administration^16,17^. The TK converts GCV into a competitive inhibitor of DNA synthesis, which leads to tumor cell lysis and the concomitant release of tumor antigens into TME^17,18^. Flt3L mediates DC recruitment and expansion in the TME, which uptake the tumor antigens, traffic them to the draining lymph nodes and prime a robust anti-tumor cytotoxic and memory T cell response, leading to tumor regression and long-term survival^9,16–18^. We first packaged TK and Flt3L into single vector of AAV (Figure 6A). It effectively reduces cell viability in vitro when treated with GCV (Figure 6B), and the infected cells secrete 600 ng/ml of Flt3L into the culture medium (Figure 6C). We then examined the delivery of AAV *in vivo* to analyze the virus distribution and Flt3l in the serum (Figure S8A). *In vivo* experiments confirmed broad viral distribution in the brain, particularly in tumor areas (Figure S8B), with markedly elevated serum Flt3L post-treatment at 40 ng/ml (Figure S8C). Importantly, mice treated with TK alone did not exhibit improved survival, confirming the necessity of the immune-modulatory effect of Flt3L (Figure S8D).

**Figure 6.**
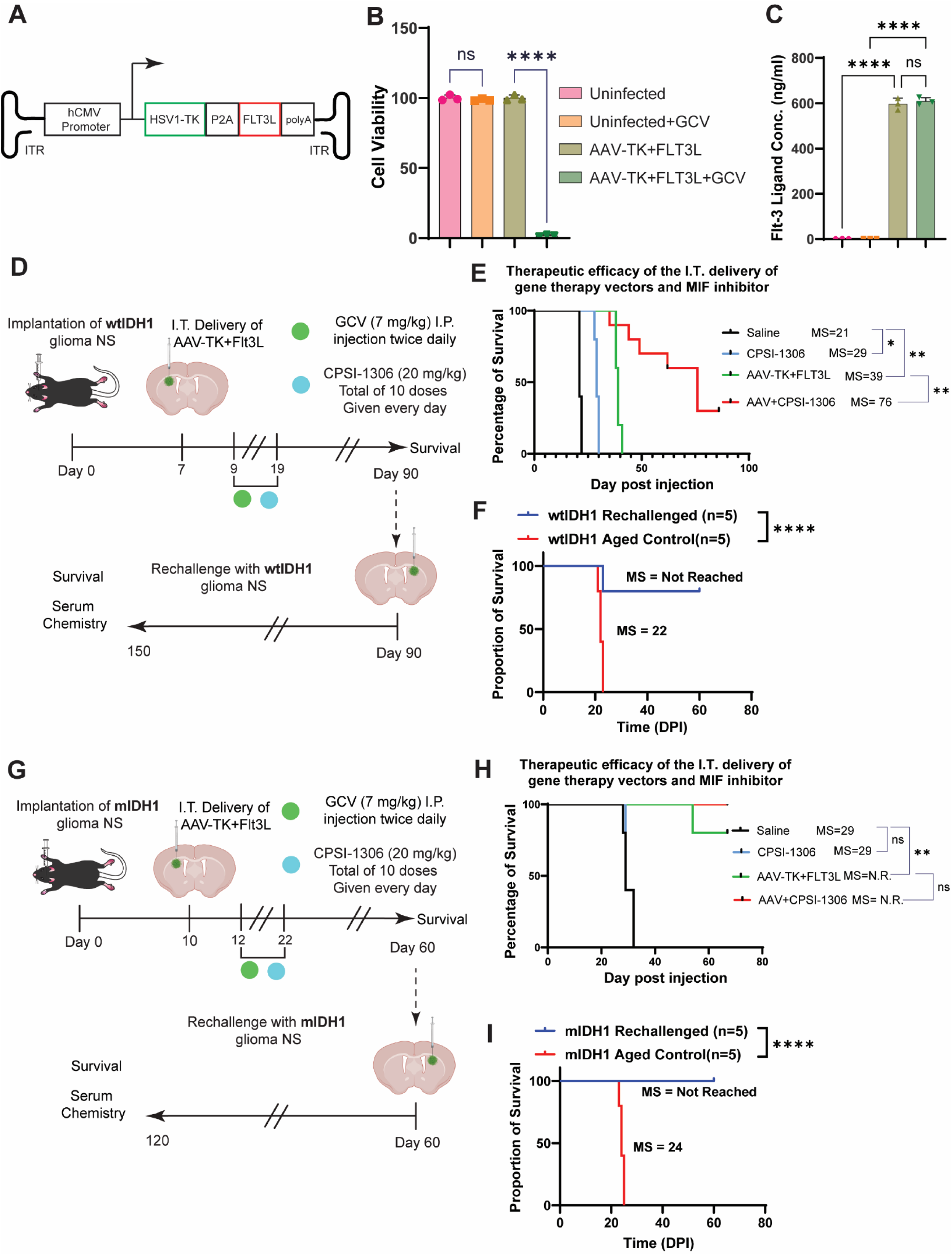
Evaluation of immune stimulated gene therapy combined with MIF inhibitor. **(A)** Illustration of AAV genome for immune stimulated gene therapy. **(B)** Proliferation of AAV and GCV treated mouse glioma cells *in vitro*. **(C)** The concentration of MIF in the conditioned media from AAV treated neurospheres was analyzed by ELISA. **(D)** Diagram depicting the experimental outline for the treatment of wtIDH1 glioma bearing mice with intratumorally delivery of AAV-TK+FLT3L gene therapy combined with MIF inhibitor. **(E)** Kaplan-Meier survival curve for the combined experiment. **** p<0.0001, Mantel-Cox test**. (F)** Kaplan-Meier survival curve for the rechallenged experiment. **** p<0.0001, Mantel-Cox test**. (G)** Diagram depicting the experimental outline for the treatment of mDIH1 glioma bearing mice with intratumorally delivery of AAV-TK+FLT3L gene therapy (GT) combined with MIF inhibitor. **(H)** Kaplan-Meier survival curve for the combined experiment. **** p<0.0001, Mantel-Cox test. **(I)** Kaplan-Meier survival curve for the rechallenged experiment. **** p<0.0001, Mantel-Cox test.

To investigate the role of MIF in immune stimulatory gene therapy, we implanted MIF knockout (MIF-KO) glioma cells into mice and administered AAV-TK/Flt3L with GCV. While either MIF ablation or gene therapy alone conferred only modest survival benefits, their combination markedly extended survival and resulted in complete tumor regression with 66.7% long term survivors (Figure S8E). Extending these findings to a translational model, wtIDH1 and mIDH1 glioma-bearing mice were treated with AAV-TK/Flt3L combined with the MIF inhibitor CPSI-1306. In wtIDH1 glioma models, combination therapy increased median survival from 39 days with gene therapy alone to 76 days, with approximately 20% of mice demonstrating long-term survival exceeding 90 days (Figure 6D, G).

Consistent with our previous results, mIDH1 glioma tumors remained responsive to gene therapy^6^; however, MIF inhibition did not alter survival (Figure 6H). These findings indicate that the MIF-CD74 axis plays a critical role in modulating therapeutic responses in wtIDH1 gliomas but is dispensable in mIDH1 tumors, where this pathway is already deficient.

To assess the development of immunological memory, long-term survivors from gene therapy experiments were rechallenged with either wtIDH1 or mIDH1 glioma cells implanted in the contralateral hemisphere, without further treatment (Figure 6D, G). As controls, naïve animals with same age were implanted with glioma cells in the striatum. Most long-term survivors from the gene therapy experiment did not develop tumors after rechallenging, whereas those from the control group succumbed due to tumor burden (Figure 6F, I). Rechallenged animals were euthanized after 60 days, perfused, and the brains were paraffin embedded and sectioned for histological analysis (Fig. S9A). After H&E staining, no tumor mass was detected in the rechallenged animals (Fig. S9A).

Also, the histological assessment of the liver of these mice did not show any abnormalities (Fig. S9B). Additionally, serum metabolites were analyzed on the rechallenged animals, showing no alterations in any marker compared to the reference range (Fig. S9C), demonstrating that the gene therapy does not elicit toxic effects. These data demonstrates that TK/Flt3L gene therapy combined with MIF inhibitors is effective for the treatment of glioma and that it prompts the development of anti-glioma immunological memory.

## Discussion

This study delineates the distinct immunological and molecular landscape of gliomas driven by mutant IDH1 (mIDH1) compared to wildtype IDH1 (wtIDH1), providing insight into the cellular mechanisms underlying tumor progression and response to immune stimulatory therapies. By leveraging single-cell RNA sequencing of human patient samples and mouse glioma models, genetic knockout models, and AAV based gene therapy approaches, we demonstrate that mIDH1-mediated epigenetic reprogramming profoundly reshapes the tumor immune microenvironment (TIME), primarily by reducing the presence of monocyte-derived tumor-associated macrophages (Mo-TAMs) and suppressing macrophage migration inhibitory factor (MIF) signaling.

MIF has been shown to play multiple roles in glioma pathogenesis^19–21^. Our analysis revealed that mIDH1 gliomas have reduced levels of MIF, driven by diminished H3K4me3 enrichment at the MIF promoter region, confirming a direct link between oncometabolite-induced epigenetic alterations and immune gene regulation. Restoration of MIF expression following treatment with IDH1 inhibitor (AG-881) in patient-derived glioma cells highlights the reversibility of these epigenetic effects and underscores the potential of therapeutically targeting mutant IDH1 to modulate the TME.

Characterization of the wtIDH1 and mIDH1 glioma TIME reinforced that myeloid cells constitute the predominant immune cell population. Notably, mIDH1 gliomas exhibited fewer Mo-TAMs and elevated Mg-TAMs relative to wtIDH1 gliomas, both in human patients and mouse models, mirroring previous reports of IDH mutation-driven shifts in myeloid lineage program and infiltration. Furthermore, our data establishes that mesenchymal-like wtIDH1 glioma cells are the main source of tumor-derived MIF, and not the host macrophages or microglia, as borne out by knockout experiments and serum ELISA analyses.

Gene knockout models elucidated the broader functional role of MIF within the TIME. Loss of MIF in glioma cells significantly enhanced survival of tumor bearing host, coinciding with a marked reduction in mature myeloid cell infiltration. Similarly, CD74 knockout in the host, or CD74 myeloid-lineage-specific deletion, resulted in extended survival and reduced macrophage and dendritic cell numbers in tumors. These findings affirm the critical role of the MIF–CD74 signaling axis in recruitment and maintenance of immunosuppressive myeloid populations, positioning it as an attractive therapeutic target.

Therapeutically, we demonstrated that combining immune-stimulatory gene therapy (TK/Flt3L) with MIF inhibition generates synergistic survival benefits in wtIDH1 glioma models. The single TK gene therapy had no effect in wtIDH1 glioma bearing mice revealed the necessity for dendritic cell recruitment via Flt3L to elicit robust anti-tumor responses. Notably, only the combination of TK/Flt3L gene therapy with MIF inhibition led to 20% long-term, tumor-free survival and the development of durable immunological memory, as demonstrated by resistance to tumor rechallenge. This combinatorial approach also proved effective using pharmacologic MIF inhibitors, indicating translational potential for clinical application.

In contrast, mIDH1 gliomas exhibited reduced baseline MIF levels and a fundamentally distinct immune microenvironment, rendering direct MIF inhibition ineffective for survival benefit while remaining responsive to immune stimulation via gene therapy alone.

These findings highlight a critical genotype-dependent therapeutic divergence: wtIDH1 gliomas require MIF pathway modulation to overcome immune suppression and maximize therapeutic efficacy, whereas mIDH1 gliomas, with their inherently deficient MIF-CD74 signaling, respond effectively to immune-stimulatory gene therapy without additional MIF targeting. This precision medicine framework suggests that IDH1 mutational status should inform treatment selection, with wtIDH1 patients potentially benefiting from MIF-targeted combination immunotherapy, while mIDH1 patients may achieve therapeutic benefit from immune-stimulatory approaches alone.

Collectively, our findings reveal crucial links between epigenetic regulation, myeloid cell heterogeneity, and immune signaling in glioma, especially in the context of IDH1 mutation status. The demonstration that MIF-CD74-driven myeloid cell recruitment shapes glioma progression provides a compelling rationale for the integration of targeted immunomodulatory strategies with gene therapy. This work offers a foundation for translational approaches combining gene therapy, small molecule inhibitors, and precision immunotherapy to improve outcomes for wtIDH1 glioma patients.

## Author Contributions

Conceptualization, Z.Z., P.R.L. and M.G.C.; Methodology, Z.Z.; Investigation, Z.Z., N.C.K., A.E.G., M.A., Y.L., A.A.M., B.L.M. and G.S. Software, J.L., J.D.W. Writing – Original Draft, Z.Z. and M.G.C.; Writing–Review & Editing, Z.Z., P.R.L. and M.G.C.; Funding Acquisition, P.R.L. and M.G.C.; Resources, W.N.A., J.H., J.D.L. and R.B.; Supervision, P.R.L. and M.G.C.

## ACKNOWLEDGMENTS

The laboratories of MGC and PRL are supported by the National Institutes of Health (NIH)/National Institute of Neurological Disorders and Stroke (NIH/NINDS) grants R37-NS094804, R01-NS105556, R01-NS122536, R01-NS124167, and R21-NS123879, and Rogel Cancer Center Scholar Award to M.G. Castro; NIH/NINDS grants R01-NS076991, R01-NS082311, R01-NS096756, R01NS122234, and National Institutes of Health/National Cancer Institute (NIH/NCI) R01-CA243916 to P.R. Lowenstein; the Department of Neurosurgery; and The Pediatric Brain Tumor Foundation, Leah’s Happy Hearts Foundation, Ian’s Friends Foundation (IFF), Chad Tough Foundation, Smiles for Sophie Forever Foundation to M.G. Castro and P.R. Lowenstein; R.B. is supported by NIH grant AR078334. We would like to acknowledge the University of Michigan Epigenomics Core, the Advance Genomics Core, the Flow Cytometry Core, and the In-Vivo Animal Core (IVAC) for providing services that contributed to this study.

## Materials and Methods

### Genetically engineered glioma mouse model

All animal studies were conducted according to guidelines approved by the IACUC of the University of Michigan (protocols PRO00011290). The wtIDH1 and mIDH1 glioma models used in this study were generated previously for our group using SB Transposon System^22^. The plasmid used to generate those models are: (i) SB transposase and LUC (pT2C-LucPGK-SB100X, henceforth referred to as SB/Luc), (ii) a constitutively active mutant of NRAS, NRAS-G12V (pT2CAG-NRASV12, henceforth referred to as NRAS), (iii) a short hairpin against p53 (pT2-shp53-GFP4, henceforth referred to as shp53), (iv) a short hairpin against ATRX (pT2-shATRX53-GFP4, henceforth referred to as shATRX), and (v) mutant IDH1^R132H^ (pKT-IDH1^R132H^-IRES-Katushka, henceforth referred to as mIDH1). Neonatal P01 C57BL/6 mice were used in all experiments. The genotype of SB-generated mice involved these combinations: (i) NRAS, shP53, and shATRX (WT-IDH1) and (ii) NRAS, shp53, shATRX, and IDH1^R132H^ (mIDH1). Mice were injected according to a previously described protocol ^23^. Plasmids were mixed in mass ratios of 1:2:2:2 or 1:2:2:2:2 (20 μg plasmid in a total of 40 μL plasmid mixture) with in vivo-jetPEI (Polyplus Transfection, 201-50G) (2.8 μL per 40 μL plasmid mixture) and dextrose (5% total) and maintained at room temperature for at least 15 min prior to injection. The lateral ventricle (1.5 mm AP, 0.7 mm lateral, and 1.5 mm deep from lambda) of neonatal mice (P01) was injected with 0.75 μL plasmid mixture (0.5 μL/min) that included: (1) SB/Luc, (2) NRAS, (3) shp53, (4) with or without shATRX, and (5) with or without IDH1^R132H^. To monitor plasmid uptake in neonatal pups, 30 μL of luciferin (30 mg/mL) was injected s.c. into each pup 24-48h after plasmid injection. In vivo bioluminescence was measured on an IVIS Spectrum (Perkin Elmer, 124262) imaging system. For the IVIS spectrum, the following settings were used: automatic exposure, large binning, and aperture f=1. For in vivo imaging of tumor formation and progression in adult mice, 100 μL of luciferin solution was injected i.p. and mice were then anesthetized with oxygen/isoflurane (1.5-2.5% isoflurane). To score luminescence, Living Image Software Version 4.3.1 (Caliper Life Sciences) was used. A region of interest (ROI) was defined as a circle over the head, and luminescence intensity was measured using the calibrated unit’s photons/s/cm2/sr. Multiple images were taken over a 25min period following injection, and maximal intensity was reported.

The strains of mice used in the study were C57BL/6 (the Jackson Laboratory, strain no. 000664), LysM-cre (the Jackson Laboratory, strain no. B6.129P2-Lyz2tm1(cre)Ifo/J, stock no. 004781), MIF-knockout mice, CD74 knockout mice and CD74 flox/flox mice were kindly provided by Dr. R. Bucala.

### Primary glioma neurosphere cultures (NS)

Mouse glioma neurosphere (NS) cultures were generated as previously described^22^. Briefly, brain tumors in mice were harvested at the time of euthanasia by intracardial perfusion with Tyrode’s solution only. The tumor mass was dissociated using non-enzymatic cell dissociation buffer, filtered through a 70 µm strainer and maintained in neural stem cell medium DMEM/F12 with L-Glutamine (Gibco, 11320-033), B-27 supplement without Vitamin A (1X) (Gibco, 12587-010), N-2 supplement (1X) (Gibco, 17502-048), Penicillin-Streptomycin (100X) (10,000 IU Penicillin) (10,000 µg/mL Streptomycin) (Corning, Cellgro, 30-001-CI), and Normocin (1X) (Invivogen, ant-nr-1) at 37°C, 5% CO_2_. Human-EGF (Peprotech, 100-15) and human-FGF (Peprotech, 100-18) were added twice weekly at 1 µL (20 ng/µL each stock) per 1 mL medium for a final concentration of 20 ng/mL. For reduced growth conditions, human-EGF and human-FGF were added twice weekly at 1 µL (20 ng/µL) per 3 mL medium for a final concentration of 6.6 ng/mL.

### Implantable syngeneic murine glioma models

Female C57BL/6 mice, aged 6-8wk old, were used for implantation models. Intracranial tumors were established by stereotactic injection of 5×10^4^ wtIDH1 or mIDH1 mouse tumor NS into the right striatum using a 22-gauge Hamilton syringe (1 μL/min) with the following coordinates: +1.00 mm anterior, 1.8 mm lateral, and 3.5 mm deep. The presence of tumors was verified 5 days post-implantation (DPI) by bioluminescence imaging.

### Human glioma cell cultures

The generation of human glioma SJ-GBM2 stably transfected cells expressing mIDH1 has been described previously^22^. SJ-GBM2 cells (CVCL_M141) were a gift of the COG Repository at the Health Science Center, Texas Tech University (Lubbock, Texas, USA). SJ-GBM2 cells were grown in IMDM (Gibco, Thermo Fisher Scientific) supplemented with 20% FBS at 37.0°C, 5% CO_2_, and were used in early passages and tested regularly for mycoplasma.

All human glioma cells derived from patients were cultured at 37.0°C with 5% CO_2_. Primary human mIDH1 glioma cultures, SF10602, from Dr. J. Costello at UCSF^24^, MGG119 from Dr. Cahill’s group at Harvard University^25^, and our laboratory generated LC1035 from a surgically resected glioma case at Umich^26^ were cultured in a tumor stem medium composed of Neurobasal-A and DMEM/F12 (1:1 ratio), further supplemented with 10 mM HEPES buffer, 1 mM MEM sodium pyruvate, 0.1 mM MEM non-essential amino acids, GlutaMAX-I (1x), B-27 without vitamin A (1x), Antibiotic-Antimycotic (1x), and Normocin (1x). EGF, FGF, PDGF-AA, and PDGF-BB were added twice a week.

### Human Single-cell RNA Sequencing

Some scRNA-seq data were obtained from Alghamri *et al.^6^* We prepared additional samples, nuclei were prepared from frozen tissue (5-25 mg) Chromium Nuclei Isolation Kit (10X Genomics) following the manual. As nuclei capture was 55%–60% less efficient than cell capture, we aimed to capture 10,000 nuclei per sample. The Chromium Single Cell 3′ (10X Genomics, Ve3) protocol was strictly followed to prepare libraries. The 10X libraries were sequenced on the Illumina NovaSeq 6000 Sequencing System. Cell Ranger V7 (10X Genomics) was used with default parameters to demultiplex reads and align sequencing reads to the genome, distinguish cells from background, intronic counts were included, and obtain gene counts per cell. Alignment was performed using the hg38 reference genome build, coupled with the Ensembl transcriptome (v110). For each sample, cells were filtered using the following quality control (QC) metrics: mitochondrial content (indicative of cell damage), number of genes, and number of unique molecular identifiers (UMIs), using the Seurat v5 ^27^. Thresholds for each sample were selected according to the distribution of each metric within the sample, which varies with sequencing coverage and the number of cells captured. UMI counts and mitochondrial content were regressed from normalized gene counts and the residuals z-scored gene-wise. Dimensionality reduction was performed using principal component analysis (PCA) applied to the most variant genes, and PCA was used as input for projection to two dimensions using uniform manifold approximation and projection (UMAP) and clustering using a shared nearest neighbor modularity optimization algorithm. Cell types were classified by reference mapping with Azimuth against GBmap reference dataset ^12^.

### Mouse Single cell-RNA sequencing

Tumors from SB mice harboring either wtIDH1 or mIDH1 were harvested and kept in media. Tumors were cut into approximately 1 mm³ pieces, transferred to a 50 ml tube, and centrifuged at 1500 rpm for 5 minutes. The pellet was resuspended in 1 ml StemPro Accutase® cell dissociation reagent (Gibco, NH) using large pore pipette tips and incubated at 37°C for 5–10 minutes to achieve complete dissociation. Accutase® was neutralized with 9 ml RPMI media and mixed by pipetting. Cells were passed through a 70 µm strainer (Alkali Inc.) attached to a 50 ml conical tube and centrifuged again at 1500 rpm for 5 minutes. Debris and dead cells were removed using the Dead Cell Removal column (Miltenyi Biotec). Cells were washed twice with PBS, counted, and resuspended at 1000 cells/µL. Viability was determined to be >90% by Countess automated cell counter. Single-cell libraries were prepared following the Chromium Single Cell 3′ protocol (10x Genomics, v3) and sequenced on the Illumina NovaSeq 6000 system. Sequencing reads were demultiplexed and aligned using Cell Ranger v7 (10x Genomics) with default parameters. CellChat analysis was performed to study cellular crosstalk between tumor cells and the tumor microenvironment ^28^.

### Cleavage Under Targets & Release Using Nuclease (CUT&RUN)

CUT&RUN was performed using CUTANA ChIC/CUT&RUN kit (Epicypher #14-1048) following manufacturer’s protocol. Briefly, 1 × 10^6^ NS cells were coupled with Concanavalin A beads, permeabilized with 0.01% Digitonin and incubated overnight at 4°C with 0.5 μg target antibody in antibody buffer (H3K4me3 Antibody - SNAP-Certified™ for CUT&RUN, Epicypher #13-0041, H3K27ac Antibody, SNAP-Certified™ for CUT&RUN and CUT&Tag, Epicypher #13-0059). The following day, cells were first incubated for 10 min with micrococcal nuclease fused to proteins A and G (pAG-MNase), which was then activated by CaCl_2_ addition. After 2 hours incubation at 4°C, the reaction was stopped with stop buffer and E. coli DNA was added as spike-in DNA. DNA was then isolated using SPRI beads and quantified with Qubit. Libraries were prepared using CUTANA™ CUT&RUN Library Prep Kit, following manufacturer’s recommendations. Fragment size was detected using a Tape Station system and multiplexed libraries were sequenced on AVITI24 sequencer (Element Biosciences) at 150bp paired end reads. The raw data quality was assessed using FastQC 0.11.3, and the adapters were trimmed using TrimGalore 0.4.4. ChIP-seq and input reads were aligned to the mouse reference genome (mm39) using bowtie2^29^ with default options, and the unique mapped reads were extracted for peak calling. The post-alignment quality control and visualized were performed by Deeptools2 3.5.1^30^.

### Western blot

Mouse NS and human glioma cells (1.0 x 10^6^ cells) were harvested for each genotype, and total protein extracts were prepared in a RIPA lysis and extraction buffer (Thermo Fisher Scientific, Pierce, 89900) with 1X of Halt protease and phosphatase inhibitor cocktail (Thermo Fisher Scientific, 78442). 40 μg of protein extract (determined by bicinchoninic acid assay (BCA), Pierce, 23227) were separated by 4-12% SDS-PAGE (Thermo Fisher Scientific, NuPAGE, NP0322BOX) and transferred to nitrocellulose membranes (Bio-Rad, 1620112). The membrane was probed with 1:1000 of a rabbit anti-MIF Ab (Abcam, ab175189), 1:2000 of an anti-β-actin Ab (Cell Signaling Technology, 3700), then followed by secondary [Dako, Agilent Technologies, goat anti-rabbit 1:4000 (P0448), rabbit anti-mouse 1:4000 (P0260)]. Enhanced chemiluminescence reagents were used to detect the signals following the manufacturer’s instructions (SuperSignal West Femto, Thermo Fisher Scientific, 34095). Blots were imaged using a ChemiDoc (Bio-Rad ChemiDoc™ MP System).

### ELISA

To assess and quantify the cytokines secreted by the wtIDH1 or mIDH1 NS, one million cells were incubated in T25 flasks for 48 hours and then the supernatants were centrifuged for 5 minutes at 500 xg to remove cell debris. Levels of MIF in the conditioned media were measured by ELISA using the manufacturer protocols (Human MIF DuoSet ELISA, DY289 or Mouse MIF DuoSet ELISA, DY1978). Briefly, to a 96-well plate 50 μl of assay diluent was added followed by 50 μl of sample, or standard; plate was incubated at 25 °C for 2 hours. Then, wells were washed five times with wash buffer and incubated for 2 h at 25 °C with 100 μl of conjugate solution. Wells were washed a further five times with wash buffer and incubated for 30 minutes at 25 °C with 100 μl of substrate solution. The reaction was stopped by adding 100 μl of stop solution to each well and the absorbance was acquired at 450 nm (background was subtracted by setting the wavelength at 540 nm).

### Flow cytometry

At symptomatic stages due to tumor burden, mice were euthanized, transcardially perfused using Tyrode’s solution, and tumors were dissected from the brain and made into single cell suspensions. The strainer was washed two times with 10 mL complete media to force cells through and to minimize cell loss. Then, tumor-infiltrating immune cells were enriched using with 30% / 70% Percoll (GE Lifesciences) density gradient. Briefly, cells were centrifuged, and the supernatant was discarded. Then, cells were resuspended in 7 mL of complete media in 15 mL Falcon tube, and 3 ml of 90% stock isotonic percoll™ (SIP, GE Healthcare) were added and mixed well by pipetting up and down 3–5 times. To layer the 70% percoll™ under the 30% percoll™ gradient, 1 mL serological pipette was filled with 1 mL of 70% percoll™ and pushed to the bottom of the 15 mL falcon tube, and the 70% percoll™ was slowly released making sure a clear interface is formed between the two gradients. Finally, the solution was spun at 800 × g for 20 min at room temperature, and the tumor-infiltrating immune cells were collected by carefully isolating the white band that formed at the interface between the two gradients.

For antibody staining, all the reactions were performed at 4°C to minimize cellular metabolic activity and marker expression changes during the staining procedure. Cells were first stained with viability dye (Alexa Fluor® 780, Fisher Scientific) to label the dead cells, after which cells were blocked with either fluorescence conjugated CD16/32 for 10 min at 4° C. Monocytic and Polymorphonuclear MDSCs were labeled with anti-CD45 (Biolegend), anti-CD11b (Biolegend), anti-Ly6G (Biolegend) and anti-Ly6C (Biolegend) antibodies (M-MDSC phenotype: CD45^+^ CD11b^+^ Ly6C^hi^ Ly6G^+^; PMN-MDSC phenotype: CD45^+^ CD11b^+^ Ly6C^lo^ Ly6G^-^). Mature dendritic cells were labeled with anti-CD45 (Biolegend), anti-CD11c (Biolegend) and anti-MHC II (Biolegend) antibodies (DC phenotype: CD45^+^ CD11c^+^ MHCII^hi^). Macrophages were labeled with anti-CD45 (Biolegend) and anti-F4/80 (Biolegend) antibodies. Total T cells were labeled with anti-CD45 (Biolegend) and anti-CD3 (Biolegend) antibodies. Antibody staining was carried out for 30 minutes, on ice, protected from light. Flow data was measured by using a BD FACSAria™ II Flow Cytometer or Cytek Aurora spectral flow cytometer (Cytek Biosciences) and analyzed using FlowJo version 10 (BD).

### In Vivo AAV Administration

The TK sequence was cloned from pAL119-TK (Addgene Plasmid #21911) and the Flt3L sequence from pAL119-Flt3L (Addgene Plasmid #21910), under the CMV promoter. The sequences were linked by P2A and insert into pAAV transfer plasmid using seamless cloning with the following primers: TK_fwd: 5’-ttgggattcgcgagaattctctagagccaccatggcttcgtacccctgc-3’; TK_rev: 5’-ccacgtcgccggcctgcttcagcagggagaagttggtggcgttagcctcccccatctc-3’; FLT3L_fwd: 5’-gctgaagcaggccggcgacgtggaggagaaccccggccccacagtgctggcgccagcc-3’; FLT3L_rev: 5’-tcacagggatgccacccgtggatcctcagggctgacactgcagc-3’. AAV-PHP.eB^31^ was packaged and titered by the University of Michigan Vector Core. Seven days after tumor implantation, AAV-PhP.eB-CMV-TK_FLT3L was delivered by intratumoral injection at a dose of 1 × 10¹¹ gc/mouse in 2 μL of PBS.

### Drug administration

An orally bioavailable small-molecule inhibitor of MIF, CPSI-1306^32^ (MedChemExpress, HY-110095), was used at a dose of 20 mg/kg/d per mouse. The mice were orally fed 100 μL per day of vehicle (10% DMSO, 40% PEG-300,5% Tween-80 and 45% Saline) or CPSI-1306 (in vehicle).

### Complete Blood Cell Counts and Serum Biochemistry

Whole blood from mice was collected into EDTA anticoagulant tubes (BD Biosciences) and complete blood counts (CBCs) were run on a Heska Element HT5 (Heska Corporation, Loveland, CO) automated veterinary hematology analyzer. Or whole blood was collected into serum separator tubes (ThermoFisher Scientific), allowed to clot, and separated into serum by centrifugation. Serum chemistries were run on an AU480 chemistry analyzer (Beckman Coulter Inc, Brea, CA). Assays were performed within the ULAM Pathology Core laboratory at the University of Michigan.

### Perfusion, fixation and paraffin embedding

After glioma NS implantation, animals were monitored daily for signs of morbidity, including ataxia, impaired mobility, hunched posture, seizures, and scruffy fur. Animals displaying symptoms of morbidity were transcardially perfused using Tyrode’s solution (0.8% NaCl (Sigma Aldrich), 0.0264% CaCl_2_2H_2_O (Sigma Aldrich), 0.005% NaH_2_PO_4_ (Sigma Aldrich), 0.1% glucose (Sigma Aldrich), 0.1% NaHCO_3_ (Sigma Aldrich), 0.02%KCl (Sigma Aldrich), followed by fixation with 4% paraformaldehyde (PFA; VWR) in Dulbecco’s modified phosphate buffered saline (DPBS; Gibco). Mouse brains were then removed from the skull and fixed in 4% PFA for an additional 48 hours at 4 °C. Then, brains were dehydrated by immersing them in solutions of increasing amounts of ethanol and lastly xylene using an automatic processor (HistoCore PELORIS 3 Premium Tissue Processing System, Leica), at the Histology Unit of the Orthopaedic Research Laboratories, at the University of Michigan. Brains were then embedded in paraffin using a Leica ASP 300 paraffin tissue processor/Tissue-Tek paraffin tissue embedding station (Leica) at the Histology Unit of the Orthopaedic Research Laboratories, at the University of Michigan.

### Hematoxylin and eosin and immunohistochemistry

Paraffin-embedded tissue was sectioned using a rotary microtome (Leica) set to 5 μm in the z-direction. Hematoxylin and eosin (H&E) staining was performed as described by us previously^23^. For immunohistochemistry (IHC), antigen retrieval slides were placed in a pressure cooker in citrate buffer (10 mM Citric Acid (Sigma Aldrich), 0.05% Tween 20 (Sigma Aldrich), pH 6). After endogenous peroxidase quenching in 0.3% H_2_O_2_ (Sigma Aldrich) and blocking in 10% goat serum in TBST for 2 hours, samples were incubated with primary antibodies (GFAP (Millipore Sigma, AB5541, 1:200), MBP (Millipore Sigma, MAB386, 1:200, CD68 (Abcam, ab125212 1:1000), and Nestin (Novus Biologicals, (NB100-1604), 1:1000)) overnight at RT. The next day, sections were washed 3 times with TBS-Tx and incubated with biotinylated secondary antibody at 1:1000 dilution in 1% goat serum TBST in the dark for 2 h. Biotin-labeled sections were developed using 3,3′-diaminobenzidine (DAB) with nickel sulfate precipitation according to the manufacturer’s instructions. Brain sections were then washed three times, followed by dehydration with xylene, and finally, coverslips were placed on the slides using DePeX Mounting Medium (Electron Microscopy Sciences). Images were obtained using brightfield/epifluorescence (Carl Zeiss MicroImaging) and analyzed using LSM5 software (Carl Zeiss MicroImaging).

### Statistics

All statistical analysis were performed using the Prism (GraphPad) software package. Data is represented as mean ± standard deviation (SD) or standard error of the mean (SEM). Statistical significance between 2 groups was determined by a student’s t-test. Statistical significance between groups of 3 or more was determined by a one-way or two-way ANOVA, followed by the Tukey’s multiple comparison test or Sidak’s multiple comparison test. Overall survival data including in animal models were plotted as Kaplan-Meier curves with Mantel-Cox test. For all tests, data were considered significant if p-values were below 0.05 (95% confidence intervals). The sample size (n) along with the statistical test performed and corresponding p-values are indicated in each figure legend. P-values are shown as follows: * p < 0.05, ** p < 0.01, *** p < 0.001, and **** p < 0.0001.

---

**Figure S1.**
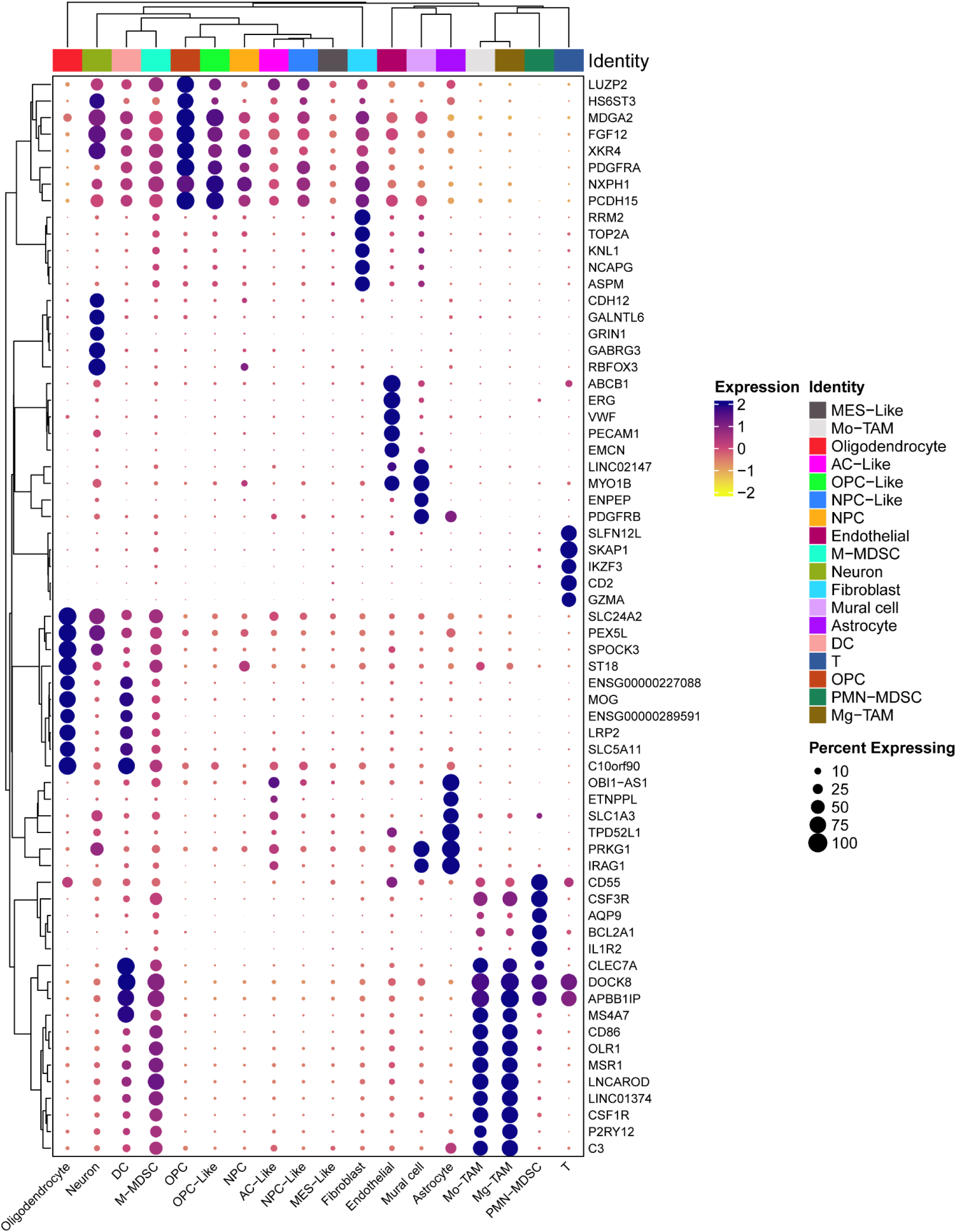
Dot plot of marker genes of human scRNA-seq, related to. Figure 1. Dot plot of top 5 markers for all populations.

**Figure S2.**
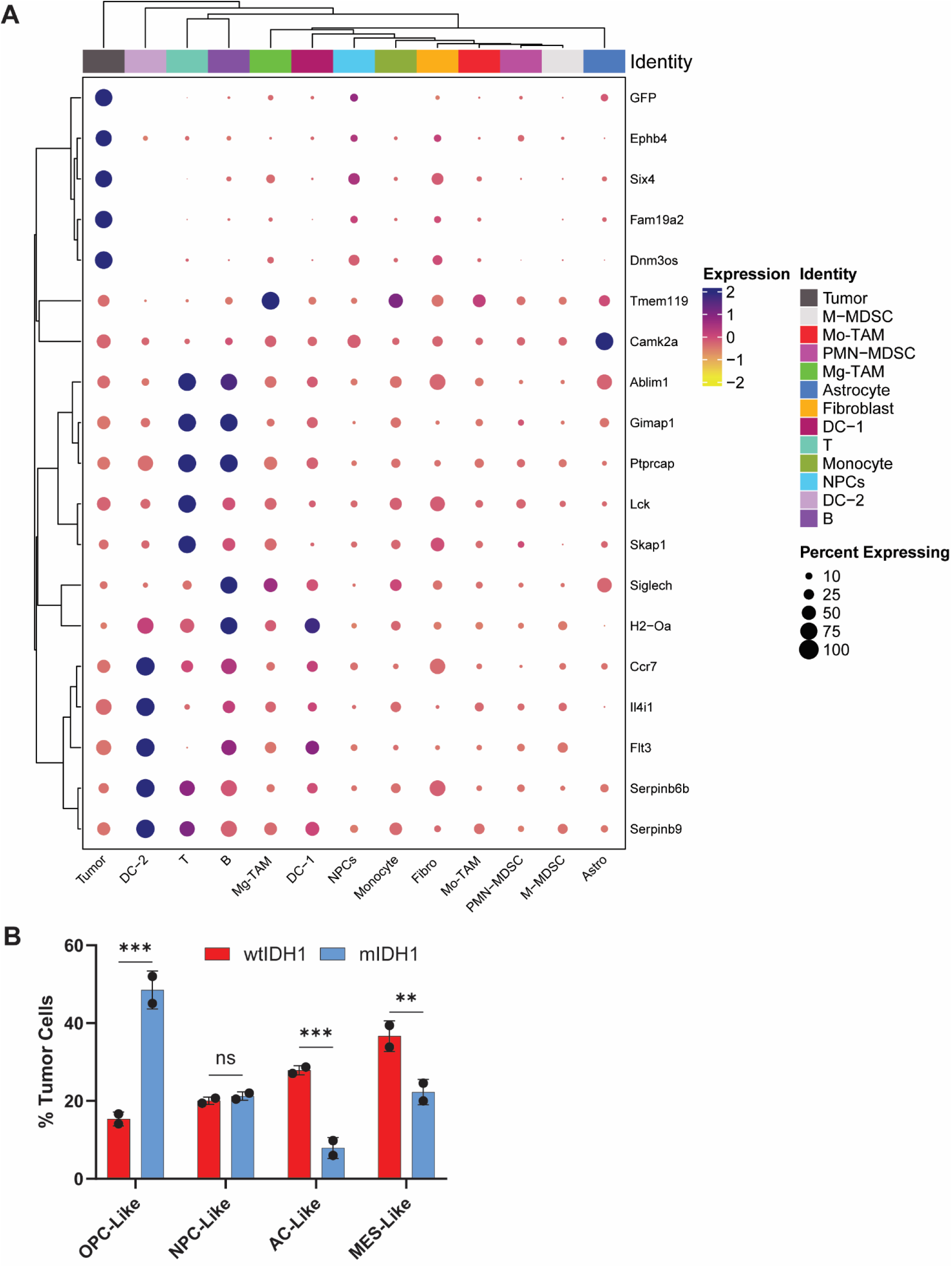
ScRNA-seq analysis for mouse wtIDH1 and mIDH1 models, related to. Figure 2**. (A)** Dot plot of top 5 markers for all populations of mouse scRNA-seq. **(B)** Four states analysis of mouse glioma cells from scRNA-seq.

**Figure S3.**
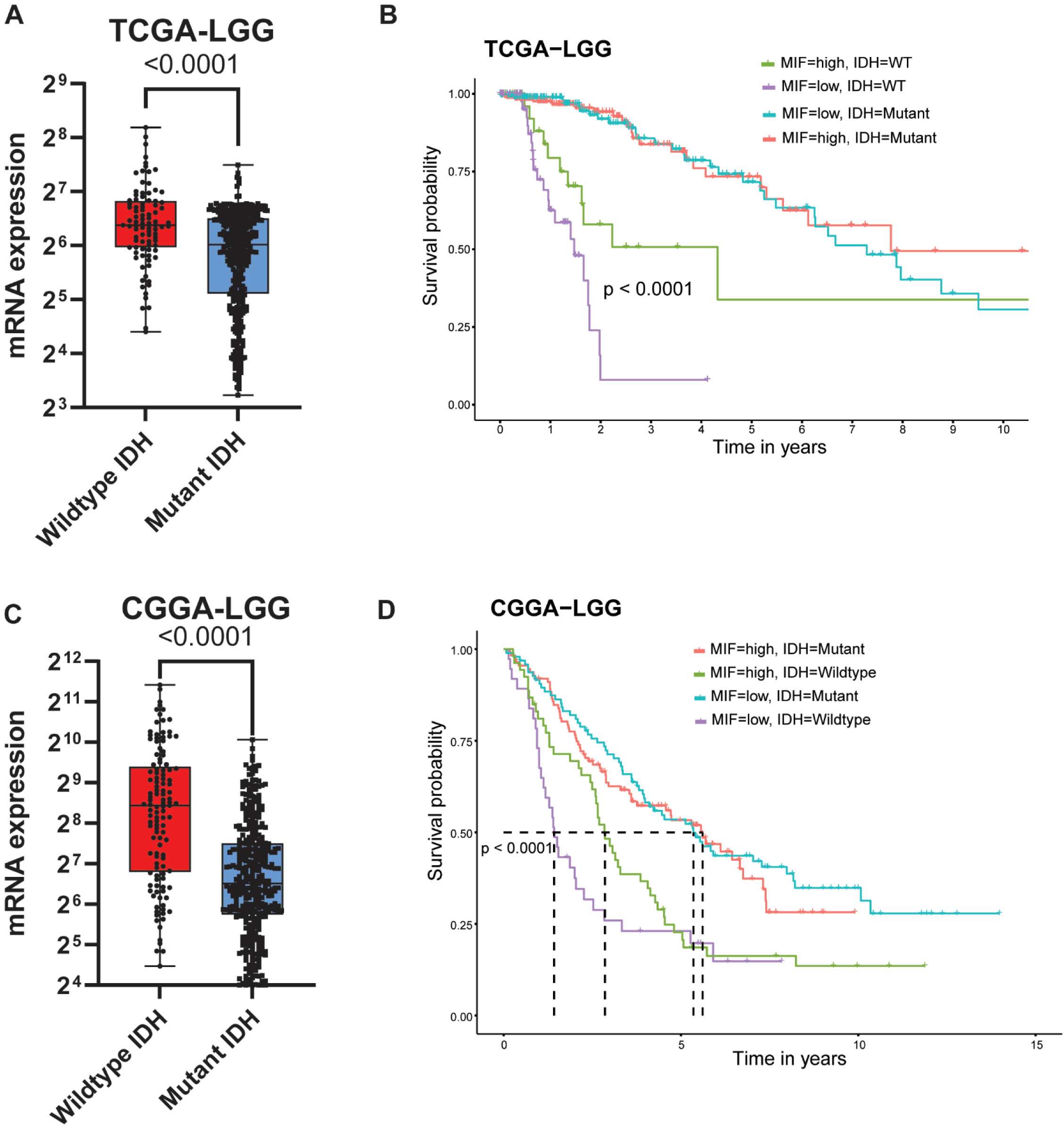
MIF is overexpressed in wtIDH1 glioma that is associated with poor clinical outcomes, related to. Figure 1**. (A)** Analysis of normalized log2 MIF mRNA expression levels in human Lower Grade Glioma (LGG) patients from The Cancer Genome Analysis (TCGA) LGG dataset segregated based on IDH mutational status as either wildtype IDH or mutant IDH. *p<0.05, unpaired t test. **(B)** Kaplan-Meier survival analysis using the log-rank test for TCGA LGG patients for whom IDH mutational status and prognosis data were available. Patients were subdivided by median expression level of MIF and IDH1 status: wtIDH1 high MIF (green), wtIDH1 low MIF (purple), mIDH1 high MIF (red) and mIDH1 low MIF (blue). ****p<0.0001; log-rank test. **(C)** Analysis of normalized log2 MIF mRNA expression levels in human Lower Grade Glioma (LGG) patients from Chinese Glioma Genome Atlas (CGGA) LGG dataset segregated based on IDH mutational status as either wildtype IDH or mutant IDH. *p<0.05, unpaired t test. **(D)** Kaplan-Meier survival analysis using the log-rank test for CGGA LGG patients for whom IDH mutational status and prognosis data were available. Patients were subdivided by median expression level of MIF and IDH1 status: wtIDH1 high MIF (green), wtIDH1 low MIF (purple), mIDH1 high MIF (red) and mIDH1 low MIF (blue). ****p<0.0001; log-rank test.

**Figure S4.**
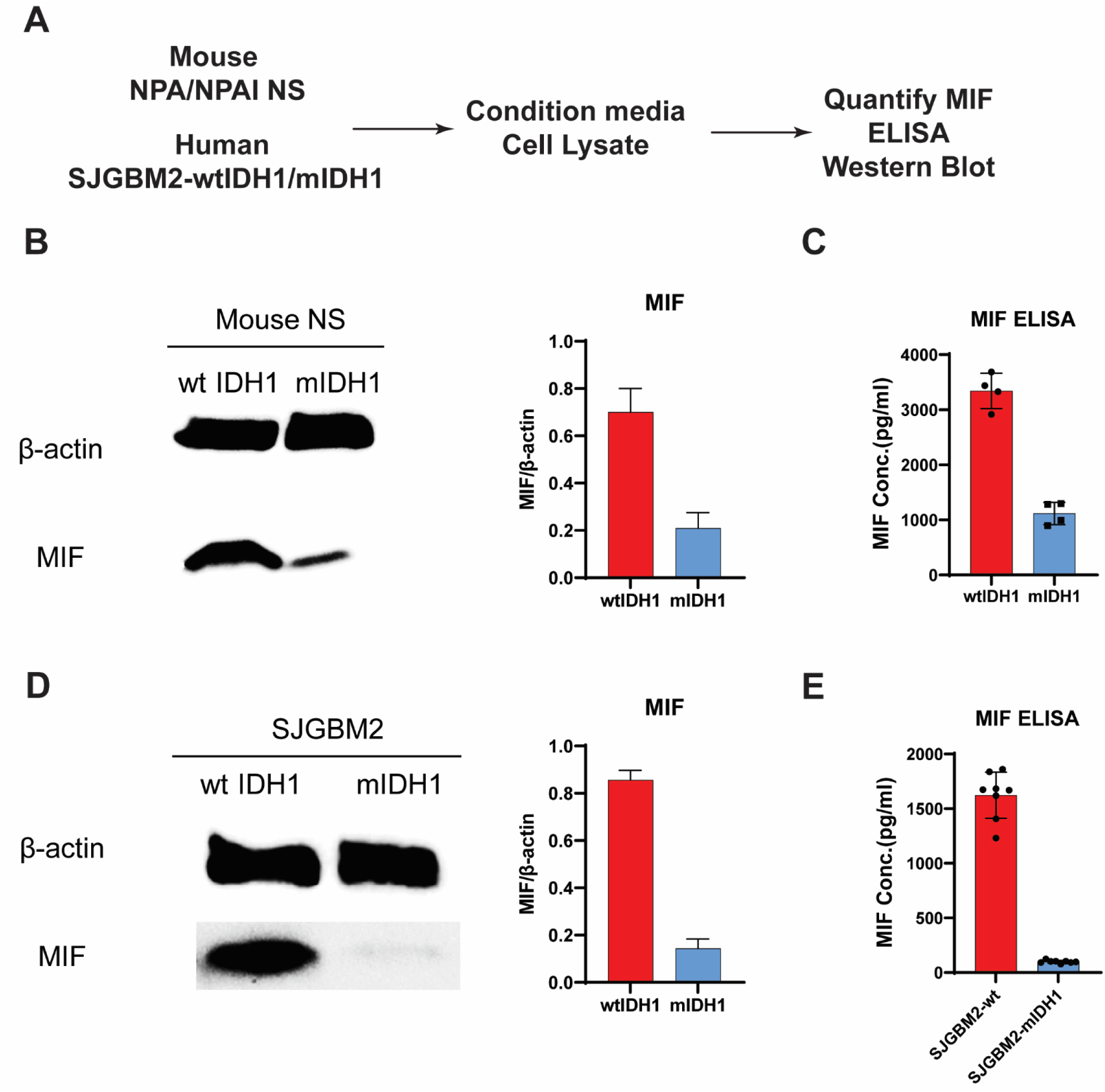
mIDH1 glioma cells exhibit lower MIF expression compared to wtIDH1 cells, related to. Figure 3**. (A)** Illustration of experimental procedure followed for the characterization of MIF expression in wtIDH1 and mIDH1 cells. **(B)** Western blot of MIF in wtIDH1 and mIDH1 mouse cells. **(C)** The concentration of MIF in the conditioned media from three different clones of wtIDH1 and mIDH1 neurospheres was analyzed by ELISA. **(D)** Western blot of MIF in wtIDH1 and mIDH1 human glioma cells. **(E)** The concentration of MIF in the human wtIDH1 and mIDH1 cell conditioned media was analyzed by ELISA.

**Figure S5.**
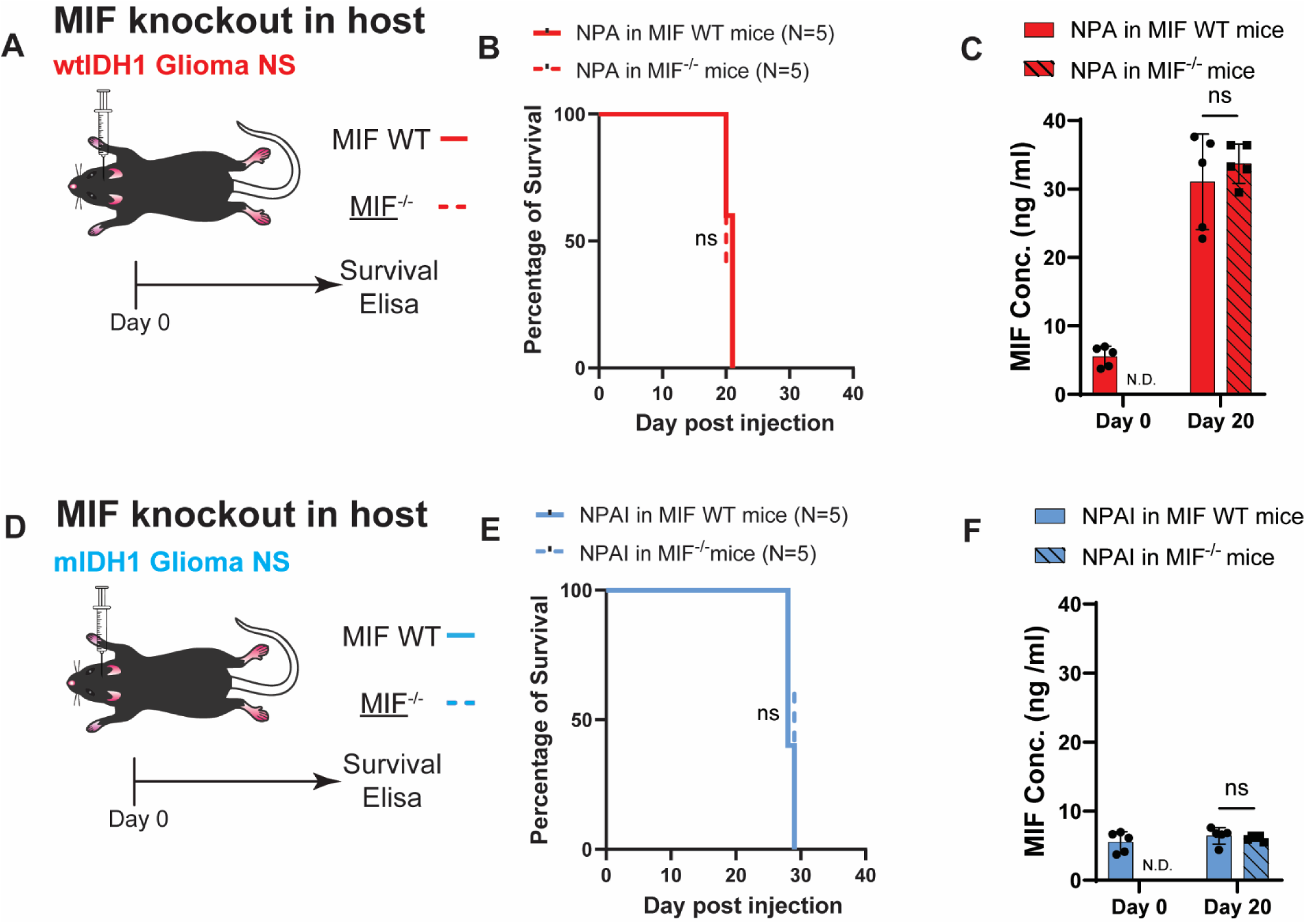
WtIDH1 glioma cells are the main source of MIF, related to. Figure 3**. (A)** Illustration of experimental procedure followed for the survival study and Elisa in MIF WT and MIF^-/-^ mice implanted with wtIDH1 mouse NS. **(B)** Kaplan-Meier survival curve shows the survival of MIF WT and MIF^-/-^ mice implanted with wtIDH1 mouse NS. Statistical significance was assessed using the Log-rank (Mantel-Cox) test. **(C)** Serum concentration of MIF analyzed by ELISA in the MIF WT and MIF^-/-^ mice implanted with wtIDH1 mouse NS. N.D., not detectable. **(D)** Illustration of experimental procedure followed for the survival study and Elisa in MIF WT and MIF^-/-^ mice implanted with mIDH1 mouse NS. **(E)** Kaplan-Meier survival curve shows the survival of MIF WT and MIF^-/-^ mice implanted with mIDH1 mouse NS. Statistical significance was assessed using the Log-rank (Mantel-Cox) test. **(F)** Serum concentration of MIF analyzed by ELISA in MIF WT and MIF^-/-^ mice implanted with mIDH1 mouse NS. N.D., not detectable.

**Figure S6.**
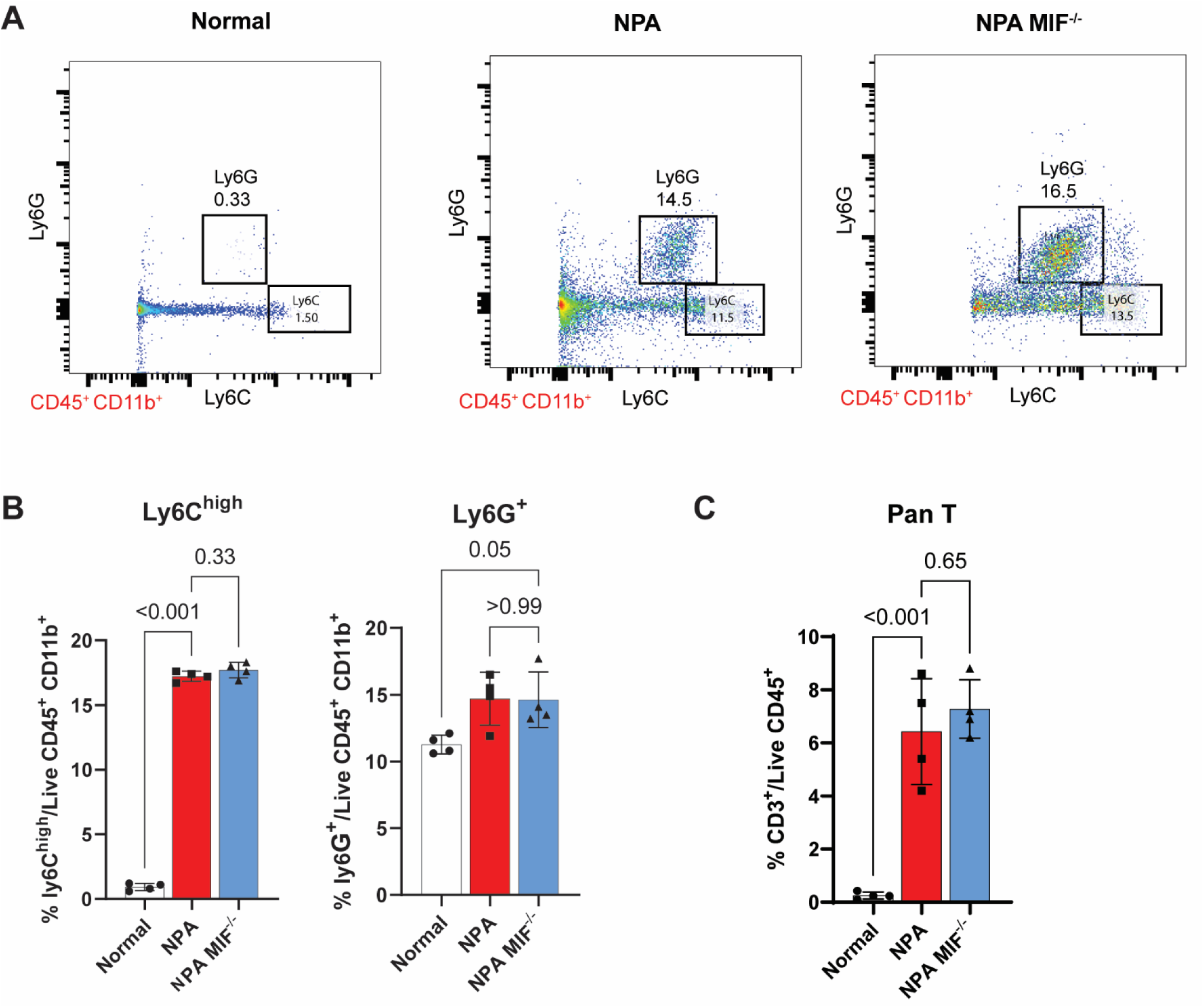
Analysis of the tumor immune microenvironment (TIME) of MIF^-/-^ glioma. (A-B) The proportion of cells with Monocytic (CD45+/CD11b+/Ly6G-Ly6Chi) or Polymorphonuclear (CD45+/CD11b+/Ly6G+Ly6Clo) myeloid derived suppressor cell. **(C)** The total proportion of pan T cells is shown as the percentage of CD45+/CD3+ cells.

**Figure S7.**
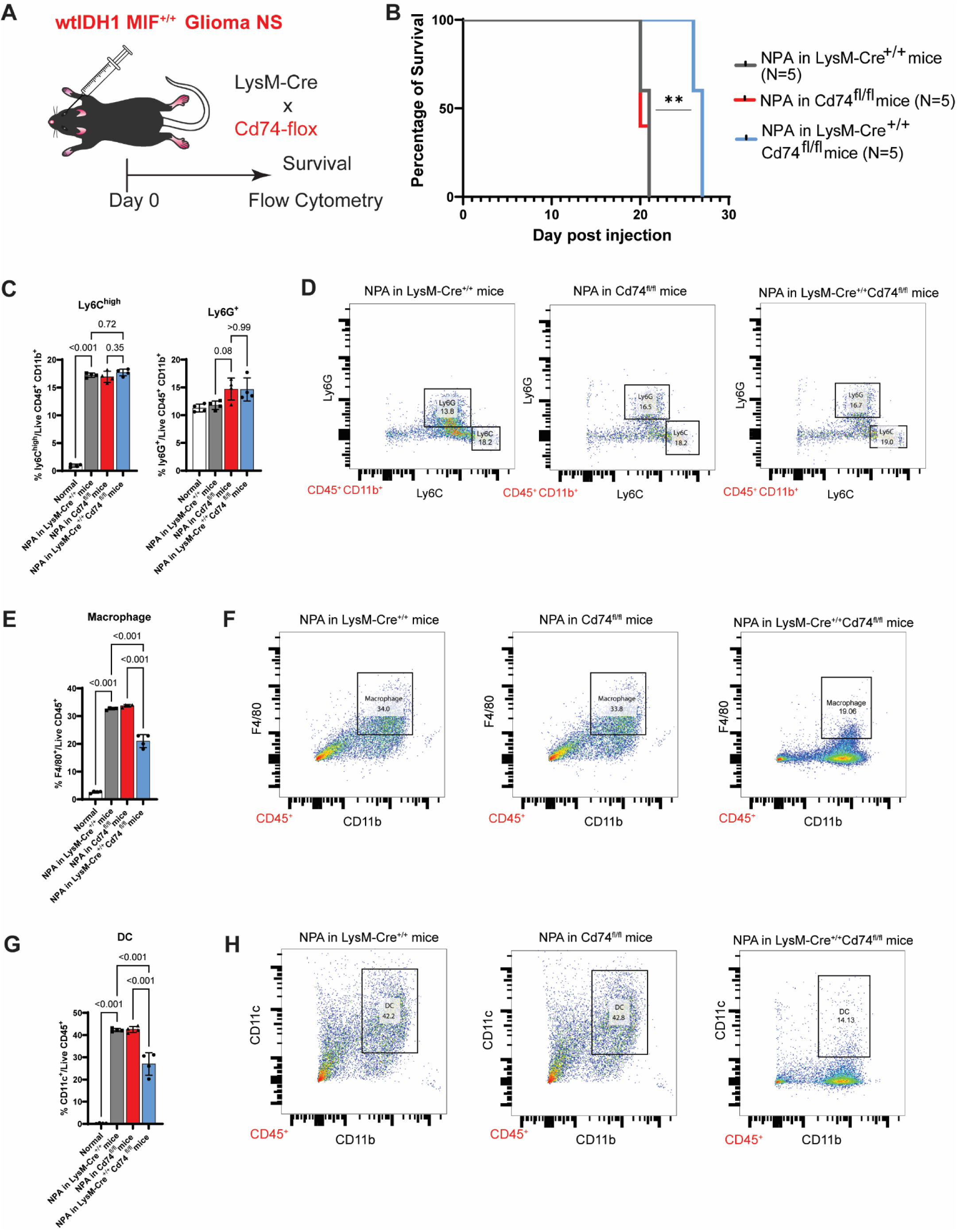
Analysis of the tumor immune microenvironment (TIME) of glioma in myeloid specific CD74^-/-^ mice. **(A)** Illustration of experimental procedure followed for the characterization of immune cells in LysM-cre, CD74^flox/flox^ and myeloid specific CD74 depleted mice implanted with WT mouse glioma cells. **(A)** Kaplan-Meier survival curve shows the survival of LysM-cre, CD74^flox/flox^ and myeloid specific CD74^-/-^ mice implanted with WT mouse glioma cells. **(C-D)** The proportion of cells with Monocytic (CD45+/CD11b+/Ly6G-Ly6Chi) or Polymorphonuclear (CD45+/CD11b+/Ly6G+Ly6Clo) myeloid derived suppressor cell. **(E-F)** The total proportion of macrophages is shown as the percentage of CD45+/CD11b+/F4/80+ cells. **(G-H)** The total proportion of DC is shown as the percentage of CD45+/CD11b+/CD11c+.

**Figure S8.**
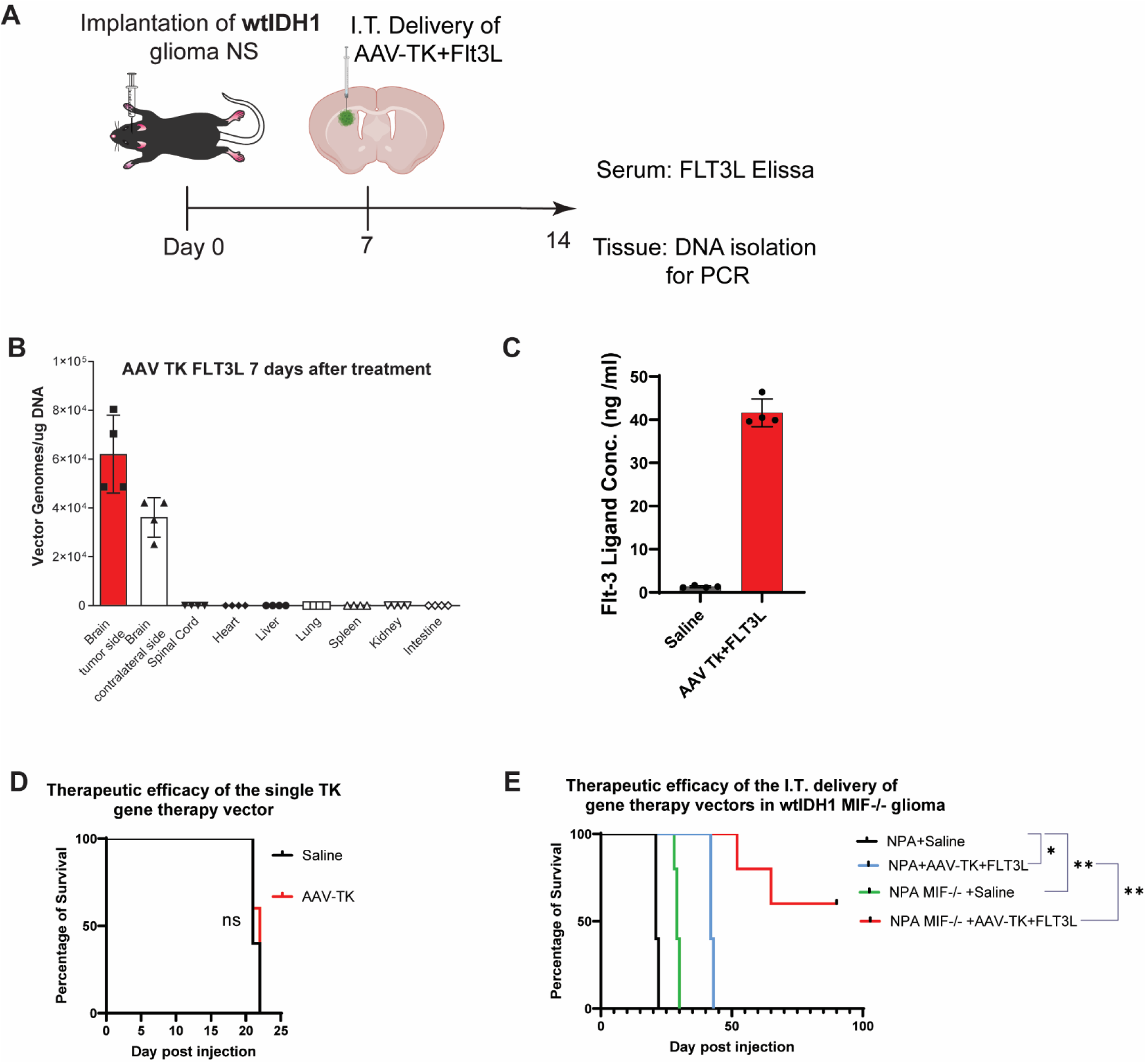
Analysis of AAV-TK+Flt3l delivery *in vivo* and evaluation of gene therapy efficacy in MIF knockout glioma bearing mice. (A) Illustration of experimental procedure followed for the characterization of AAV delivery **in vivo** after 1 week. **(B)** Quantitative PCR show the distribution of AAV in different locations. **(C)** The concentration of Flt-3 ligand in the mouse serum after 1 week was analyzed by ELISA. **(D)** Kaplan-Meier survival curves show that administration of TK alone does not provide a survival benefit, underscoring the necessity of both Ad-TK+Flt3L gene therapy vectors for a robust anti-tumor response. **(E)** Kaplan-Meier survival curves curve of NPA MIF WT and NPA MIF^-/-^glioma NS treated with or without AAV-TK+Flt3L GT.

**Figure S9.**
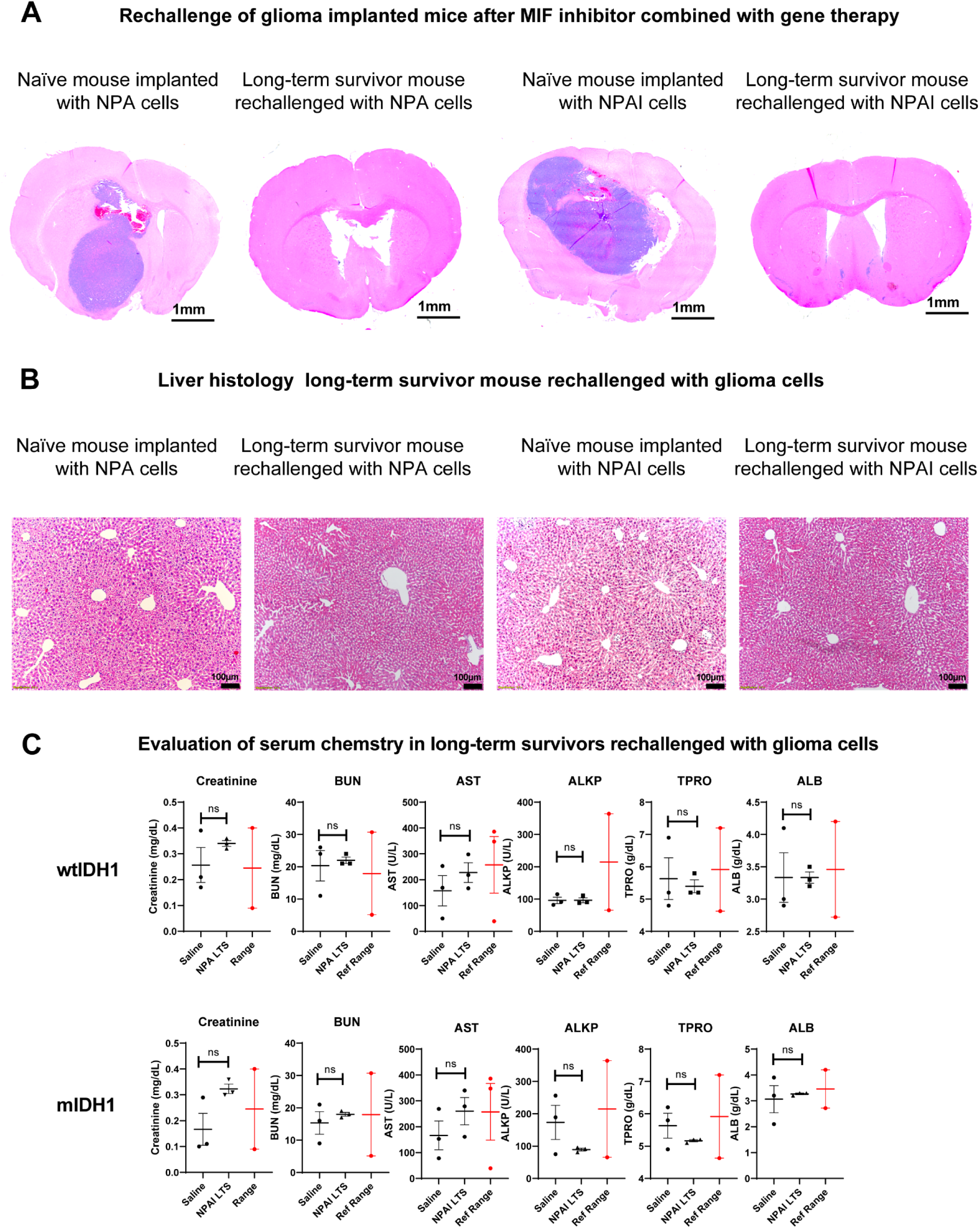
Rechallenge of long-term survivors, related to Figure 7. **(A)** Brain hematoxylin/Eosin histology from Naïve mouse implanted with wtIDH1 or mIDH1 cells versus gene therapy long-term survivor rechallenged with wtIDH1 or mIDH1 glioma cells. Long term survivors (LTS) do not show signs of tumor development due to development of immunological antitumoral memory. Black scale bar: 1 mm. **(B)** Long term survivors from combination treatments were implanted with 50,000 neurospheres in the contralateral hemisphere and 60 days post rechallenge animals were euthanized and processed for histological analysis. Paraffin embedded 5 µm liver sections were stained for hematoxylin and eosin (H&E). Black scale bar: 100 µm. **(C)** Assessment of serum metabolites in long term survivors rechallenged with glioma cells. BUN= Blood Urea Nitrogen, ALT = alanine transaminase, AST = Aspartate Transferase (AST), ALKP = alkaline phosphatase (ALP), GLUC = glucose, TPRO = total protein, ALB2 = albumin, TBIL = Total Bilirubin, Ca = Calcium. Levels are compared to a reference range (Ref. Range).

